# From Lotka-Volterra Dynamics to Community Assembly: Theory, Topography, and Empirical Applications

**DOI:** 10.64898/2026.07.14.738515

**Authors:** Sebastian J. Schreiber, Joe Brennan, Jürg Spaak

## Abstract

(1) Community assembly graphs (CAGs) summarize which species combinations can coexist and how single-species invasions drive transitions between them, encoding the pathways, alternative endpoints, and cycles that make up a community’s assembly history. Constructing CAGs from dynamical models requires methods that are both computationally tractable and faithful to the underlying ecological dynamics. However, existing methods rely on restrictive assumptions, such as global stability, that exclude alternative stable states and non-equilibrium dynamics known to occur in empirical systems.
(2) We develop a computational pipeline that constructs CAGs from any generalized Lotka–Volterra model. Building on the invasion graph framework and its connection to permanence, the pipeline verifies that community dynamics are bounded, identifies which subsets of species coexist in the sense of permanence, determines which single-species invasions are dynamically realized, and assigns each community a topographic height equal to the length of the longest assembly path leading to it. We also provide a numerical algorithm to simulate the dynamics of community assembly.
(3) We prove several general properties of the resulting graphs, including that a successful invader is never excluded by the residents it invades, and that, in the absence of assembly cycles, permanent communities can be reassembled by introducing their species one at a time in the right order. We prove that the CAG faithfully reproduces the compositional shifts seen in the numerically simulated dynamics of assembly. Applying the pipeline to three empirically based models (a New Zealand grassland, a European pasture, and a Puerto Rican ant community), we show how competition strength and mutualistic feedbacks reshape the assembly landscape and how intransitive competition generates assembly cycles.
(4) Our approach accommodates alternative stable states and non-equilibrium dynamics without requiring global stability, and it turns the long-standing landscape metaphor into a quantitative, mechanistically grounded object by resolving what “height” means. More broadly, it makes the topography of the assembly pathways measurable, providing a way to compare the historical contingency and predictability of the assembly in ecological systems.

## Introduction

Ecological communities are not static assemblages but rather the product of ongoing processes of colonization, establishment, and local extinction. Understanding how these processes give rise to observed patterns of biodiversity, and how they might be redirected by conservation and restoration interventions, is one of the central challenges of community ecology [HilleRisLambers et al., 2012, Suding et al., 2024]. The urgency of this challenge has only grown as the acceleration of global change drives rapid community reorganization through changes in resource availability, abiotic stress, and environmental variability [Suding et al., 2024]. Predicting and guiding these trajectories requires a theoretical framework capable of capturing the full complexity of community assembly: which species combinations can coexist, how invasions alter community composition, and when history matters.

The community assembly graph (CAG) has emerged as a powerful conceptual tool for meeting this need, bringing coexisting species combinations, the invasions that connect them, and the contingencies of assembly history into a single representation. In a CAG, vertices represent communities and directed edges represent transitions between communities driven by species invasions. Each vertex corresponds to a subset of the species pool that persists together in the absence of further invasions, and each edge records the successful establishment of a species arriving from outside the resident community. Assembly then becomes a directed walk through this graph, beginning at the empty community and proceeding invasion by invasion. Read as a whole, the graph encodes all possible assembly histories for a given species pool, making explicit the pathways from no species to persistent communities, the alternative endpoints that arise from priority effects, and the ongoing cycles in species composition that can prevent assembly from ever settling [Law and Blackford, 1992, Law and Morton, 1993, 1996, Schreiber and Rittenhouse, 2004, Song, 2025]. As Song [2025] has argued, the CAG functions as a kind of Rosetta Stone for assembly theory: a shared language in which informational, dynamical and probabilistic approaches can be expressed and compared.

The CAG has a long history, with foundational contributions from Post and Pimm [1983], Drake [1988, 1990, 1991], and Law and Morton [1993, 1996]. It has recently attracted renewed interest driven by theoretical advances connecting assembly to coexistence theory and network ecology [Song et al., 2021, Serván and Allesina, 2021, Hofbauer and Schreiber, 2022, Spaak and Schreiber, 2023, Song, 2025] and by growing empirical evidence for priority effects and alternative stable states [Fukami, 2015, Suding et al., 2024]. Furthermore, there is growing recognition that CAGs provide a natural framework for applied questions in restoration and conservation [Godoy et al., 2024, Suding et al., 2024]. This renewed interest has sharpened the need for methods that connect CAG structure directly to the mechanistic parameters (interaction strengths, intrinsic growth rates, functional traits) that empirical ecologists measure and manipulate.

There are two broad approaches to the construction of CAGs. The first builds CAGs from fixed structural or statistical rules, bypassing explicit population dynamics [Luh and Pimm, 1993, Song et al., 2021]. These static approaches efficiently characterize large numbers of assembly outcomes and have yielded important insights into how graph topology relates to the predictability of assembly. Luh and Pimm [1993] compared three recipes for generating transition graphs: random, landscape, and minimally constrained. They showed that even removing the simplest ecologically implausible cycles qualitatively transforms assembly behavior, with persistent cycles becoming rare, multiple persistent states becoming more common, and assembly trajectories shortening. Song et al. [2021] took a complementary approach, exhaustively enumerating assembly graphs for communities of three species and showing that four topological features predict an information-theoretic measure of community predictability: the number of alternative stable states and transient paths, the length of the cycle and the presence of escapes from cycles to stable states. Although these approaches have yielded valuable insights, they share a fundamental limitation. Because graphs are constructed without reference to population dynamics, it remains unclear which graphs are biologically realizable for any given set of species interactions, and graph properties cannot be connected to the mechanistic parameters that empirical ecologists measure.

The second approach derives CAGs directly from population models, most commonly the generalized Lotka-Volterra models. The foundational contribution here is Law and Morton [1996], who developed the first reliable and computationally tractable method for constructing CAGs from Lotka-Volterra dynamics. Their algorithm equated coexistence with permanence [Jansen and Sigmund, 1998, Hofbauer and Sigmund, 1998, Schreiber, 2000, 2006], which guarantees that all species persist despite perturbations and applies equally to equilibrium and non-equilibrium dynamics. Using this algorithm, they established that assembly sequences terminate at uninvadable or cyclic endpoints, that the number of such endpoints is small even for large species pools, and that some endpoints cannot be reassembled from their constituent species alone. More recent dynamical work has pursued CAG construction under the stronger assumption that the dynamics restricted to any subset of species converges to a globally stable equilibrium. Under this assumption, Serván and Allesina [2021] identified when community attractors correspond to end states of assembly, Godoy et al. [2024] constructed empirical CAGs for Mediterranean annual plant communities under varying environmental conditions, and Calleja-Solanas et al. [2026] classified the frequency and structure of assembly cycles. However, the assumption of global stable equilibria is ecologically restrictive. It precludes alternative stable states and priority effects [Hofbauer and Sigmund, 1998], which are well-documented features of natural communities [Fukami, 2015, Suding et al., 2024], and excludes non-equilibrium dynamics despite the fact that Lotka-Volterra models can exhibit arbitrarily complex dynamics [Gilpin, 1979, Zeeman, 1993, van den Driessche and Zeeman, 2004, Vano et al., 2006, Vandermeer and Perfecto, 2023].

A recent advance that addresses these limitations is the development of the invasion graph and its connection to permanence [Hofbauer and Schreiber, 2022, Spaak and Schreiber, 2023, Schreiber, 2025]. Like CAGs, vertices in invasion graphs represent communities, but directed edges represent *potential* transitions driven by single-*or multiple* species invasions. One can compute invasion graphs for Lotka-Volterra systems by solving systems of linear equations for the equilibria and evaluating the per-capita growth rates of absent species, i.e. invasion growth rates. Hofbauer and Schreiber [2022] proved that permanence requires that every subcommunity can be invaded by at least one missing species, and that this condition is also sufficient when the invasion graph is acyclic. These advances provide a foundation for the construction of CAGs without the restrictive assumption of global stability. However, several technical challenges must be overcome before translating this foundation into a practical pipeline for community assembly. First, community dynamics must be bounded, a condition that none of the earlier methods explicitly verifies but that must hold for invasion graphs to be useful. Although boundedness typically occurs in ecological models rooted in competitive and predator-prey interactions, it can be violated for models that account for positive interactions [May and McLean, 2007]. Surprisingly, there are publications in which this condition has not been satisfied without the authors recognizing this issue [Spaak et al., 2021]. Second, not all transitions predicted by invasion growth rates are realized by the underlying Lotka-Volterra dynamics, and one must identify and remove such false positives. Together, these challenges mean that a practical computational pipeline is needed to construct CAGs from the invasion graph framework.

Here, we address this gap by developing a tractable computational pipeline for constructing CAGs. Building on the invasion graph framework, our approach does not require global stability and therefore accommodates alternative stable states and non-equilibrium dynamics. It goes beyond invasion graphs by verifying boundedness and which transitions are dynamically realized. The pipeline also yields a dynamically grounded realization of the metaphor of the assembly landscape of Pimm [1991] and Luh and Pimm [1993]. Community states were assigned heights arbitrarily in these works, a weakness Pimm acknowledged: “what the abstraction ‘akin to height’ is, I do not know” [Pimm, 1991, page 251]. Our approach resolves this by assigning each community an elevation equal to the length of the longest directed path leading to it from the empty community, a quantity fully determined by the interaction parameters of the underlying Lotka-Volterra system rather than arbitrarily assigned. We illustrate the pipeline with two empirically parameterized Lotka-Volterra models examining how the strength of interspecific competition and mutualistic interactions shape the topography of community assembly. These examples are motivated in part by the topological categories of dominance control, facilitative control, and positive feedback control that Suding et al. [2024] identify as characteristic interaction structures of plant communities. We also analyze an empirically inspired Lotka-Volterra model with intransitive competitive dynamics that generate cycles in the invasion graph, illustrating the broader scope of the pipeline beyond the acyclic case.

## Methods

Our goal is to construct community assembly graphs that are both computable from an empirical parameterization and faithful to the dynamics they summarize. In a community assembly graph, vertices represent sets of species that coexist in the sense of permanence (defined below) and directed edges represent single-species invasions that carry one such community to another (Figure 1, top panels). Building this object from a Lotka– Volterra model requires answering four questions in turn: do the community dynamics remain bounded, which subsets of species coexist, which single-species invasions are actually realized by the dynamics, and what is the topography of the resulting assembly landscape? The sections below address each question, and together they define a six-step computational pipeline that takes an interaction matrix and growth-rate vector as input and returns the community assembly graph together with a topographic summary of assembly complexity. Because a graph is only useful if it reflects the underlying dynamics, we provide a numerical algorithm for simulating assembly. We also establish several mathematical properties of the resulting graphs, chief among them a proof that the graphs our pipeline produces correctly predict the compositional changes those simulations exhibit.

**Figure 1.**
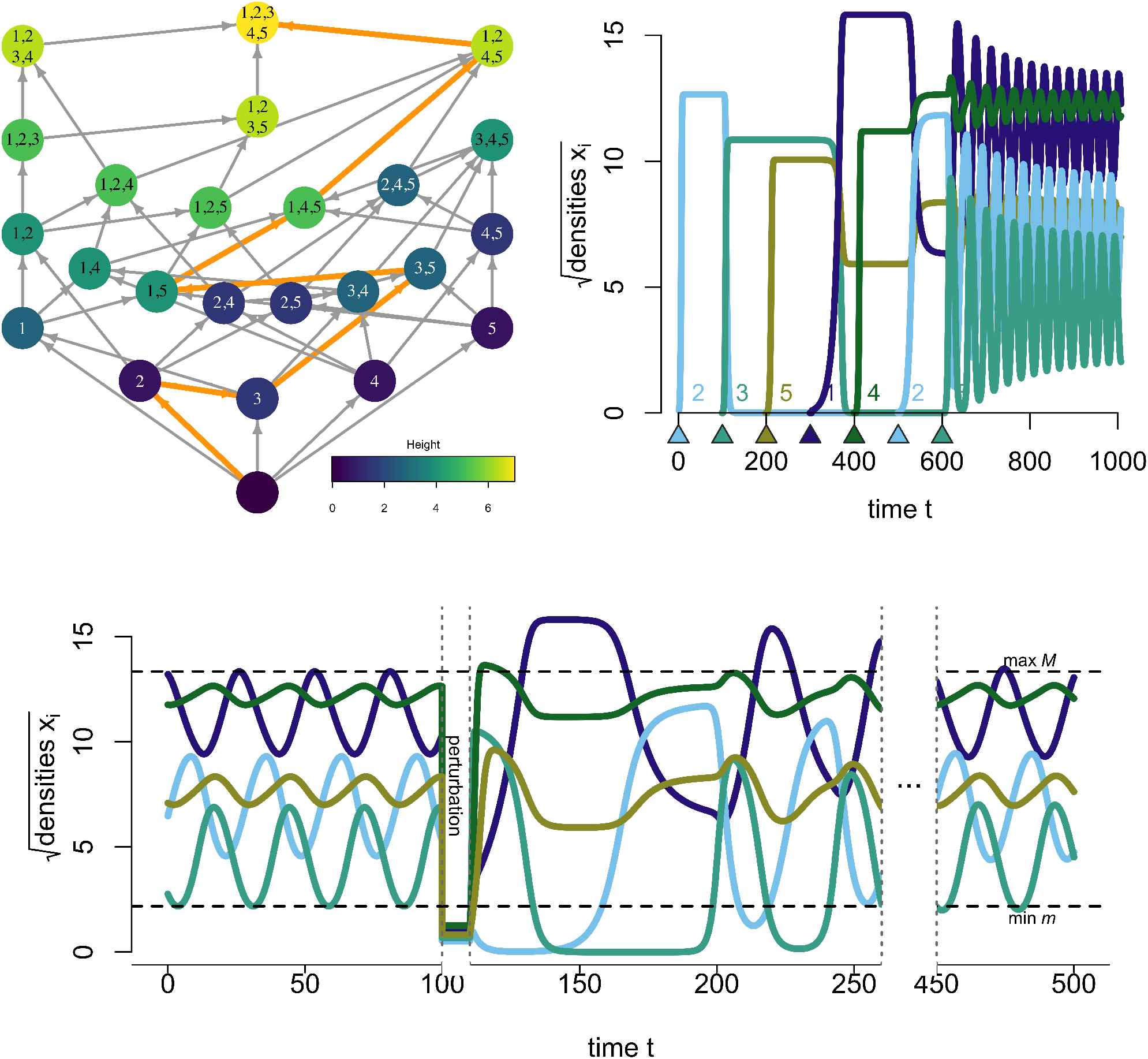
Community assembly graphs and permanence. *Top left:* A community assembly graph in which vertices represent permanent communities and directed edges represent dynamically realized single-species invasions. An assembly path to the full community is highlighted in orange. *Top right:* Simulated Lotka–Volterra dynamics in which species are introduced sequentially along the highlighted path; community composition at each stage matches the assembly graph predictions. Triangles and labels indicate the timing of species invasions. *Bottom:* Following a large perturbation (dashed vertical line), all species recover to their long-run attractor. Dashed horizontal lines mark the permanence bounds *m* and *M*, within which all species densities ultimately remain.

### Lotka-Volterra Models, Boundedness, and Permanence

To model species interactions, we focus on Lotka-Volterra models, which form the theoretical foundation for much of community ecology. In their general form, there are *k* species with densities *N*_1_, *N*_2_, …, *N*_*k*_. Each species *i* has an intrinsic per-capita growth rate *b*_*i*_, corresponding to its per-capita growth rate in the absence of all interactions. Interactions with species *j*, including conspecific interactions (*j* = *i*), have a per-capita effect *a*_*ij*_*N*_*j*_ on species *i* where the interaction coefficient *a*_*ij*_ represents the per-capita impact of species *j* on species *i*. Under these assumptions, the generalized Lotka-Volterra equations are:

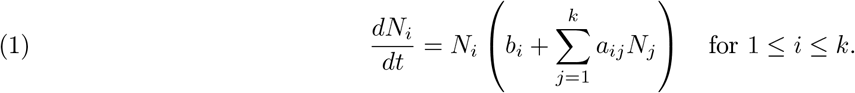

The dynamics of (1) are bounded (more formally, dissipative) if there exists a maximal density *M >* 0 such that all species densities eventually fall and remain below *M* for any initial condition *N* (0) = (*N*_1_(0), *N*_2_(0), …, *N*_*k*_(0)), i.e., *N*_*i*_(*t*) ≤ *M* for all *i* and all sufficiently large *t*. Any realistic model of community dynamics should exhibit bounded dynamics. When an interaction matrix *A* ensures bounded dynamics for any choice of the intrinsic growth vector *b, A* is known as a *B*-matrix [Hofbauer and Sigmund, 1998]. In our pipeline, we introduce a simple inductive algorithm to verify whether *A* is a *B*-matrix or not. Violations of the *B*-matrix condition arise from positive feedback loops in the interaction structure, such as pairs of mutualistic species or more complex indirect facilitation networks, which can lead to unbounded growth.

A positive equilibrium, also known as a feasible equilibrium [Saavedra et al., 2017], for a set of species *S* ⊂ {1, …, *k*} is an equilibrium 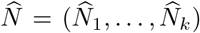 where 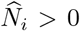 if *i* ∈ *S* and 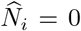 otherwise. We denote this equilibrium 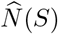 when it exists. Although the existence of a positive equilibrium is necessary for stable coexistence in a Lotka-Volterra model [Hofbauer and Sigmund, 1998], it is not sufficient: Lotka-Volterra systems can exhibit stable chaotic or periodic dynamics [Gilpin, 1979, Zeeman, 1993, Hofbauer and So, 1994, van den Driessche and Zeeman, 2004, Bobieński and Żołądek, 2005, Gyllenberg et al., 2006]. A form of stable coexistence that accommodates such nonequilibrium dynamics is permanence [Jansen and Sigmund, 1998, Hofbauer and Sigmund, 1998, Schreiber, 2000]. A community *S* is permanent if its dynamics are bounded and there exists a minimal density *m >* 0 such that, whenever all species in *S* initially have a positive density, all species densities in *S* eventually exceed and remain above *m* (Figure 1, bottom panel). Permanence ensures that all species recover to this minimal density after arbitrarily large, rare perturbations [Jansen and Sigmund, 1998].

### Invasion Growth Rates and Invasion Graphs

Permanence can be identified using invasion growth rates, also known as growth rates when rare or low density growth rates. These growth rates are the key metric of modern coexistence theory [Chesson, 1994, Ellner et al., 2019, Grainger et al., 2019]. Invasion growth rates correspond to the per-capita growth rates of rare species while the rest of the community is at a stationary state. If the species in *S* are at equilibrium 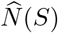, then the invasion growth rate of a missing species *i* ∈*/ S* is

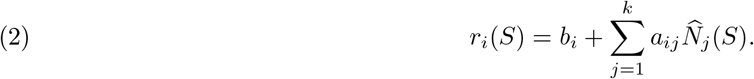

If *r*_*i*_(*S*) *>* 0, species *i* introduced at low density will initially increase exponentially; if *r*_*i*_(*S*) *<* 0, it will decrease exponentially toward extinction. A remarkable property of the Lotka-Volterra models is that invasion growth rates remain meaningful even when the resident community *S* does not settle to a fixed equilibrium. The time-averaging property [Hofbauer and Sigmund, 1998] guarantees that for any bounded solution *N* (*t*) in which all species in *S* persist and all species outside *S* are absent, the long-term per-capita growth rate of a missing species *i* equals *r*_*i*_(*S*):

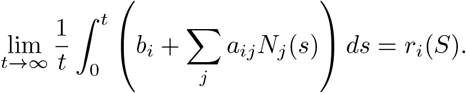

Because the interior equilibrium 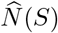 of the subsystem restricted to *S* is unique whenever the principal submatrix *A*_*S*_ is nonsingular, *r*_*i*_(*S*) is the same for every such solution. Hence invasion growth rates, and with them the invasion graph and the permanence criteria introduced below, do not depend on whether *S* settles to an equilibrium, a periodic orbit, or a chaotic attractor, nor on which attractor a solution approaches when *S* supports several. At no point does the pipeline require computing these attractors. Henceforth, we assume that all invasion growth rates are nonzero, a condition that holds generically (e.g., for randomly chosen *A* and *b*) but excludes the degenerate case of neutral coexistence.

Using equilibria and invasion growth rates, Hofbauer and Schreiber [2022] introduced the invasion graph to characterize coexistence in the sense of permanence. Unlike the community assembly graph, this graph identifies *potential* transitions between communities arising from invasions by one *or more* species. As we discuss below, the CAG is a subgraph of the invasion graph where certain edges and vertices are pruned. However, to verify permanence for any subset of species, the full invasion graph is required. The vertices of the invasion graph are all equilibria of the Lotka-Volterra model, including the extinction equilibrium. Each equilibrium is labeled with the set *S* ⊆ {1, 2, …, *k*} of species at positive density. Two distinct vertices *S* and *T* are joined by a directed edge if (i) all species *i* ∈ *T* \ *S* can invade *S*, i.e., *r*_*i*_(*S*) *>* 0, and (ii) all species *j* ∈ *S* \ *T* cannot invade *T*, i.e., *r*_*j*_(*T* ) *<* 0. The first condition requires that all species gained in the transition *S* → *T* grow when rare introduced into community *S*. The second condition requires that all species lost in the transition cannot reinvade. A key result of Hofbauer and Schreiber [2022] establishes the dynamical relevance of these edges: if there exists a community trajectory whose composition converges to an attractor associated with *T* in forward time and to an attractor associated with *S* in backward time (a *connecting orbit*, defined precisely in Supplement A), then the edge *S* → *T* must appear in the invasion graph. Hence, directed edges are *necessary* for the existence of such heteroclinic connections but not *sufficient* : the invasion graph may contain edges that are not realized by any community trajectory, i.e. false positives. The CAG removes these false positives as we will show below.

Using invasion graphs, Hofbauer and Schreiber [2022] proved necessary and sufficient conditions for permanence of the entire community {1, …, *k*}. If the community is permanent, then every subcommunity can be invaded by at least one missing species: at each equilibrium 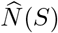with *S*≠ {1, …, *k*}, there is some *i* ∈*/ S* with *r*_*i*_(*S*) *>* 0. This invadability is *necessary* for permanence: if instead some subcommunity resisted invasion by all of its missing species, then there would be community trajectories, starting from the full species pool, that converge to that subcommunity, and the excluded species would be lost. Invadability is not, however, always *sufficient*, and whether it guarantees permanence depends on whether the invasion graph is acyclic or cyclic. A directed cycle in the invasion graph is a directed path that returns to its starting subcommunity *S*, that is, a sequence of invasions, by single or multiple species, that eventually restores the original composition. When the invasion graph is cyclic (i.e., has at least one directed cycle), invadability is necessary but is no longer sufficient: an invadable community can still fail to be permanent. In that case, there is no general characterization of permanence, though a sufficient condition of Hofbauer [1981] can be used to resolve individual cases [Spaak and Schreiber, 2023]. We note that a directed cycle in the invasion graph is a combinatorial object and should be distinguished from cycles in the dynamics. Cycles in the dynamics come in two forms with different relationships to the graph. A heteroclinic cycle, in which trajectories pass successively near a sequence of boundary equilibria as in the rock-paper-scissors dynamics of May and Leonard [1975], always produces a cycle in the invasion graph, since each of its connecting orbits yields an edge. By contrast, periodic orbits and chaotic attractors within a community leave no trace in the graph: the time-averaging property reduces them to the interior equilibrium of that community.

### The Community Assembly Graph Pipeline

Community assembly graphs describe transitions between permanent communities driven by sequential single-species invasions, each followed by sufficient time for the community to settle to a new attractor. They therefore correspond to a subgraph of the invasion graph, restricted to ecologically realizable transitions. To construct them in a mathematically rigorous fashion, we introduce a six-step computational pipeline (Figure 2). The pipeline checks whether the Lotka-Volterra model has bounded dynamics, identifies which subsets of species coexist in the sense of permanence, determines the dynamically realized single-species transitions among those communities, and characterizes the resulting assembly landscape through a height function that captures the complexity of assembly paths.

**Figure 2.**
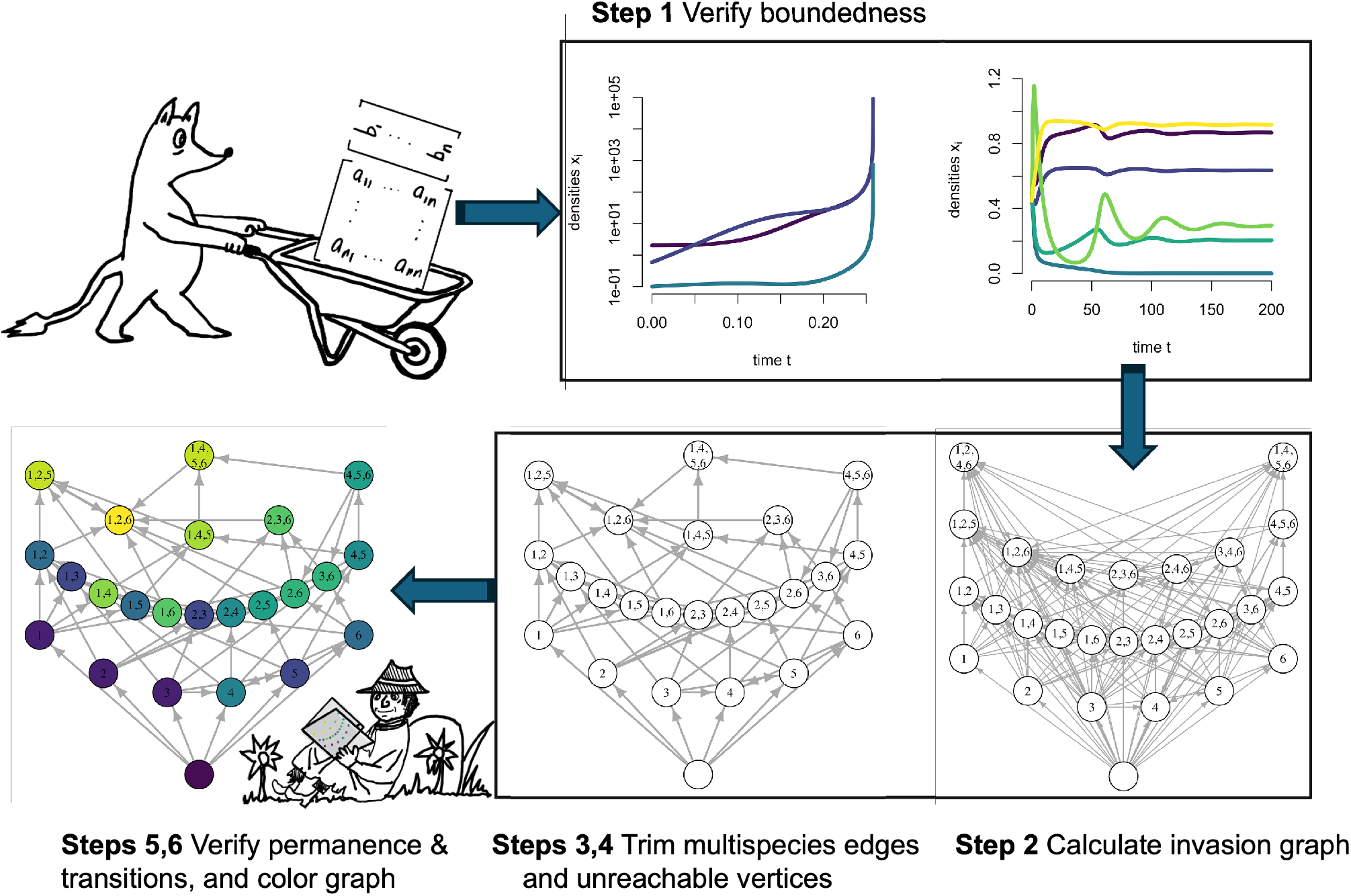
The community assembly graph pipeline. The pipeline takes an interaction matrix *A* and intrinsic growth rate vector *b* and applies six steps. Step 1 uses the *B*-matrix criterion to verify that the dynamics are bounded. If they are, Step 2 computes the invasion graph. Steps 3 and 4 prune away edges that do not correspond to single-species invasions and vertices corresponding to “garden of Eden” communities. Step 5 verifies that vertices are permanent and that edges are dynamically realized. Step 6 assigns topographic heights.

#### Step 1: Verify bounded dynamics

We provide an inductive algorithm to verify whether the interaction matrix *A* is a *B*-matrix (Supplement A). The algorithm exhaustively examines all principal submatrices of *A*: for each subset *S* ⊆ {1, …, *k*} of size *j* = 2, 3, …, *k*, it extracts the principal submatrix *A*_*S*_ = *A*[*S, S*] and computes its inverse 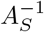. A violation of the *B*-matrix condition occurs whenever all entries of 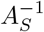 are strictly positive, i.e. 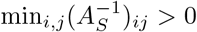. The algorithm also issues a warning if any diagonal entry *a*_*ii*_ is non-negative, since this precludes intraspecific self-regulation. *A* is a *B*-matrix if and only if no principal submatrix has an entirely positive inverse.

#### Step 2: Compute the invasion graph

The invasion graph is computed as detailed in Hofbauer and Schreiber [2022]. Briefly, all non-negative equilibria 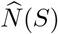are found by solving 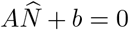 restricted to each species subset *S*, invasion growth rates *r*_*i*_(*S*) are evaluated at each equilibrium, and the invasion graph is constructed from its definition. Two conditions can prevent completion of this step: a principal submatrix of *A* may be singular, in which case equilibria are non-unique or non-existent for the corresponding species subset; or some invasion growth rates may be exactly zero, violating the genericity assumption stated earlier. In either case, the algorithm issues a warning.

#### Step 3: Identify single-species transitions

Each directed edge in the invasion graph is examined to determine whether it corresponds to an invasion by a single species. As shown in Supplement A, a directed edge *S* → *T* corresponds to a single-species invasion by species *i*, denoted 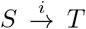, only if *r*_*i*_(*S*) *>* 0 (species *i* can invade) and *r*_*j*_(*T* ) *<* 0 for all *j* ∈ *S* ∪ {*i*} \ *T* (none of the missing species including possibly species *i* can invade *T* ). Edges that do not satisfy these conditions are removed. We note that even when *r*_i_ (*S*) *>* 0, there does not need to exist a community *T* such that 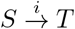; this failure is only possible when the invasion graph restricted to *S* ∪ {*i*} is cyclic (Supplement A).

#### Step 4: Remove “garden of Eden” communities

Vertices corresponding to communities that cannot be reached by sequential single-species invasions from the empty community are removed. Formally, a community *S* is removed if there is no directed path 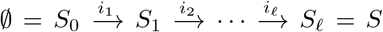 in the graph from Step 3. When the invasion graph is acyclic, the removed communities are always non-permanent communities (Supplement A).

#### Step 5: Verify permanence and realize edges

For each remaining vertex, permanence of the corresponding community is verified using the invasion graph criterion described above. When the invasion graph restricted to a vertex is cyclic, this criterion is only necessary, but permanence can be established instead using a criterion of Hofbauer [1981], which requires solving a linear programming problem (Supplement B). Each vertex is thereby classified as permanent (the invasion graph criterion holds and the restricted invasion graph is acyclic, or the Hofbauer criterion holds), not permanent (some subcommunity cannot be invaded by any species of the community that it lacks), or undetermined (the restricted invasion graph is cyclic and the Hofbauer criterion fails), and the pipeline reports this classification together with the criterion that settled it. Each remaining directed edge 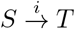 is then checked to confirm that it is dynamically realized: introducing species *i* at sufficiently low density into the resident community *S*, at any positive densities and with all other species absent, leads to the community *T* . Formally, for any *η >* 0 there exist *ξ >* 0 and *m >* 0 such that, whenever *N*_*j*_(0) ≥ *η* for all *j* ∈ *S*, 0 *< N*_*i*_(0) ≤ *ξ*, and *N*_*j*_(0) = 0 for all *j* ∈*/ S* ∪ {*i*}, one has lim inf_*t*→∞_ *N*_*j*_(*t*) ≥ *m* for all *j* ∈ *T* and lim_*t*→∞_ *N*_*j*_(*t*) = 0 for all *j* ∈ (*S* ∪ {*i*}) \ *T* . (Supplement A). A sufficient analytical condition for its realization is that (i) the invasion graph restricted to *S* ∪ {*i*} is acyclic, and (ii) *T* is the unique community satisfying *r*_*j*_(*T* ) *<* 0 for all *j* ∈ *S* ∪ {*i*} \ *T* (Supplement A). If this condition isn’t satisfied, one can run a simulation using our numerical algorithm to identify whether or not the edge is realized. If the edge is not realized, prune it from the community assembly graph and return to step 4.

The sixth and final step assigns a topography to the assembly landscape. For acyclic assembly graphs, height increases strictly along every assembly path, providing a natural measure of assembly complexity. For cyclic graphs, height is constant within each strongly connected component and increases between components.

#### Step 6: Assign topographic heights

For an acyclic community assembly graph, the height *h*(*S*) of community *S* is defined as the length of the longest simple directed path from ∅ to *S*. For cyclic community assembly graphs, the graph is first condensed by collapsing each strongly connected component to a single vertex using Tarjan’s algorithm [Sedgewick and Wayne, 2011]; the resulting condensation is acyclic, and heights are assigned to components as before, with all communities within a component receiving the same height. For a community in a strongly connected component, height therefore measures the length of the longest assembly path leading into the component, not the length of paths within it, which can be made arbitrarily long by cycling. This height assignment works because if 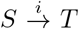, then augmenting any path of length *h*(*S*) from ∅ to *S* by the transition 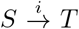 yields a path of length *h*(*S*) + 1 from ∅ to *T*, so *h*(*T* ) ≥ *h*(*S*) + 1.

When every vertex of an acyclic community assembly graph is permanent, each invasion along an assembly path adds exactly one species, the invader, and removes zero or more residents (see **Mathematical Properties of the Community Assembly Graph**). Hence, for any assembly path of length *l* from ∅ to *S*, the difference *l* −|*S*| equals the total number of resident species lost along the path, counted with multiplicity: an invasion that displaces two residents contributes two to this count. In particular, *h*(*S*) ≥ |*S*|, and the excess *h*(*S*) − |*S*| equals the number of species lost along a longest path to *S*. It is bounded below by the number of displacing invasions along that path, with equality when each such invasion removes a single resident. When *S* can be assembled by a path of length |*S*|, as often is the case (see **Mathematical Properties of the Community Assembly Graph**), *h*(*S*) − |*S*| is the range of assembly path lengths to *S*: the difference between the most direct and the most circuitous assembly paths. This range lies between two extremes. It equals zero exactly when no assembly path to *S* involves a displacement, as occurs for every community when interspecific competition is weak and all invasions succeed. At the other extreme, since a simple path visits each vertex at most once, *h*(*S*) ≤ *V* − 1, where *V* is the number of vertices in the community assembly graph. Therefore, the range is at most *V* − 1 − |*S*|. A strict competitive hierarchy of *n* species attains this bound: the graph consists of the empty community and the *n* single-species communities, and introducing species from the most to the least competitively inferior yields a path of length *n* = *V* − 1 to the community of the dominant. We use *h*(*S*) − |*S*| as a summary statistic for the complexity of the assembly landscape. Other summaries, such as the mean path length, require a probability distribution on assembly paths, for instance by weighting all paths equally or by specifying a stochastic process for the order in which species are introduced. Different choices induce different distributions of path lengths and hence different means, whereas the range *h*(*S*) − |*S*| is independent of these choices. We therefore focus on the range.

### Numerically Simulating Assembly Dynamics

To simulate the assembly dynamics of the Lotka-Volterra models, we consider a sequence of introductions of *l* ≥ 1 species {*i*_1_, *i*_2_, …, *i*_*l*_}. Each species in the sequence is introduced by adding a small density *δ >* 0 to its current density, which is zero when the species is absent from the community. Then the dynamics are numerically integrated for *τ* time steps. At the end of the numerical integration, we remove (i.e., set densities to zero) species whose densities fall below a quasi-extinction threshold *ε >* 0. The next species is then introduced, and the algorithm continues until all species in the sequence were introduced and the community dynamics were integrated for *τ* time steps. Consistent with the assumption of rare and small introductions, we choose *δ >* 0 to be small and *τ >* 0 to be large. For simplicity, we assume that quasi-extinction threshold *ε* equals the introduction density *δ*. We use these simulations to compare the predicted dynamics from the community assembly graph with the actual Lotka-Volterra dynamics.

### Mathematical Properties of the Community Assembly Graph

The community assembly graphs produced by our pipeline satisfy several analytical properties that hold regardless of the specific parameterization. These properties clarify the relationship between the assembly graph and the underlying community dynamics and provide useful guarantees for interpreting our applications. These properties are proven in Supplement A.

### Invasion implies persistence of the invader

If *S* is a permanent community in the community assembly graph, then any successful single-species invasion by species *j* ∈*/ S* results in a new community that retains the invader: 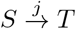 implies *j* ∈ *T* . A species that successfully invades a community cannot subsequently be excluded by the residents it just colonized, i.e., there is no “resident strikes back” phenomenon [Case, 1995, Mylius and Diekmann, 2001]. Indeed, if the invader were excluded after a successful invasion, the resulting community *T* would not contain *j*, and no missing species would be able to invade *T*, giving each missing species a negative invasion growth rate. But then *S* would violate the necessary condition for permanence, since permanence requires that at least one missing species have a positive invasion growth rate at every subcommunity. However, when coexistence is instead equated with the existence of a locally stable equilibrium, Case [1995] showed that the “resident strikes back” phenomenon can occur in Lotka–Volterra models.

### Minimal assembly of permanent communities

If *S* is permanent and the invasion graph restricted to *S* is acyclic, then *S* can be reached from the empty community by single-species invasions and *S* is not removed in Step 4. *S* remains in the community assembly graph unless Step 5 prunes an edge among its subcommunities, which happens in none of our examples. Hence, when the invasion graph is acyclic, every community removed in Step 4 is non-permanent. Intuitively, permanence of *S* implies that each subcommunity of *S* is invadable by another species in *S*, and acyclicity ensures that each invasion leads to a new community. Therefore, one can construct a sequence of single-species invasions from species in *S* that must end in a noninvadable community, which is *S* itself. Only intransitive loops, such as rock-paper-scissors, prevent the assembly of permanent communities through single-species introductions. Furthermore, if every vertex in the community assembly graph is permanent and Step 5 prunes no edges, then each community *S* can be assembled by |*S*| species introductions: there is an ordering of the species in *S* such that introducing them one at a time, in this order, assembles *S*. Thus, every such community is reachable by the shortest possible assembly sequence, one in which each invasion succeeds and adds a species, so that richness increases at every step.

### Community assembly graphs capture the dynamics of assembly

Provided that all of the vertices of the community assembly graph are permanent and all the transitions satisfy the dynamical realization check in Step 5 of the pipeline, we provide a mathematical proof in Supplement A that all paths in the community assembly graph correctly predict the changes in community composition for the numerical algorithm for simulating assembly dynamics whenever introductions are sufficiently small and sufficiently rare, i.e. *δ >* 0 is sufficiently small and *τ >* 0 is sufficiently large.

### End states are dynamical attractors

If *S* is a permanent end state of the assembly graph, then *S* is locally attracting for the full Lotka–Volterra dynamics: for any initial condition with all species in *S* at positive density and all remaining species at sufficiently low density (i.e., *N*_*i*_ *>* 0 for all *i* ∈ *S* and *N*_*j*_ ≪ min_*i*∈*S*_ *N*_*i*_ for *j* ∈*/ S*), the community composition converges to an attractor associated with *S*. When the invasion graph is acyclic and *S* is the unique end state, the guarantee is stronger: *S* is a global attractor, and the composition of the community converges to *S* from any initial condition with all species present. This result confirms that end states of the assembly graph are not merely graph-theoretic absorbing states, but genuine long-run outcomes of the community dynamics.

## Application to Empirical Lotka -Volterra models

To illustrate the scope and utility of our methods, we apply the community assembly graph framework to two empirically parameterized Lotka–Volterra models and one empirically inspired Lotka-Volterra model. In addition to illustrating how the pipeline works, the two empirical examples address the question: how does the strength of interspecific competition and facilitation shape the topography of community assembly? We characterize this topography through four complementary lenses: the size of the community assembly graph, measured by its numbers of vertices and edges; the overall rate of invasion success, the fraction of missing species able to invade feasible communities; the number and species richness of end states; and the excess height *h*(*S*) − |*S*| of end states, which measures how circuitous assembly to them can be. The empirically inspired example illustrates how the methods apply to invasion graphs containing cycles. The examples span a range of community types and data sources and are chosen to complement rather than duplicate one another: a New Zealand grassland [Roxburgh, 1994, Roxburgh and Wilson, 2000] in which interspecific competition is systematically varied; an Irish managed pasture [Geijzendorffer et al., 2011] in which reciprocal facilitation between two species is varied; and a Puerto Rican coffee agroecosystem [Vandermeer and Perfecto, 2023] whose intransitive competitive dynamics generate cycles in the invasion graph.

### Impacts of the strength of interspecific competition on the assembly of a New Zealand grassland community

For our first empirical application, we use a data set from Roxburgh [1994] and Dormann [2008] involving a 7 species grassland community. The species (*Agrostis capillaris, Festuca rubra, Hydrocotyle heteromeria, Holcus lanatus, Prunella vulgaris, Ranunculus repens*, and *Trifolium repens*) were grown in monoculture and in all pairwise combinations under an additive design, with aboveground biomass harvested after one year. Carrying capacities were estimated directly from monoculture yields, and pairwise competition coefficients were estimated by fitting a nonlinear function to biomass data in replicates, producing a full 7 × 7 competition matrix *A* and an intrinsic growth rate vector *b* for use in a Lotka–Volterra community model.

As all entries in the interaction matrix *A* are negative, the *B*-matrix condition is satisfied immediately. Computing the invasion graph (upper left panel in Fig. 3) reveals that it is acyclic. Although the invasion graph has 46 vertices corresponding to the empty community and 45 positive equilibria for non-empty communities, 15 of these vertices are not reachable by single-species introductions and, consequently, excluded from the community assembly graph (upper right panel in Fig. 3). As the invasion graph is acyclic, our mathematical results imply that all 15 of these removed vertices correspond to non-permanent communities. The remaining vertices are all permanent. Moreover, all edges in the community assembly graph satisfy the dynamical realization condition. Therefore, the community assembly graph is mathematically guaranteed to correctly capture the numerically simulated dynamics of assembly (lower panels in Fig. 3).

**Figure 3.**
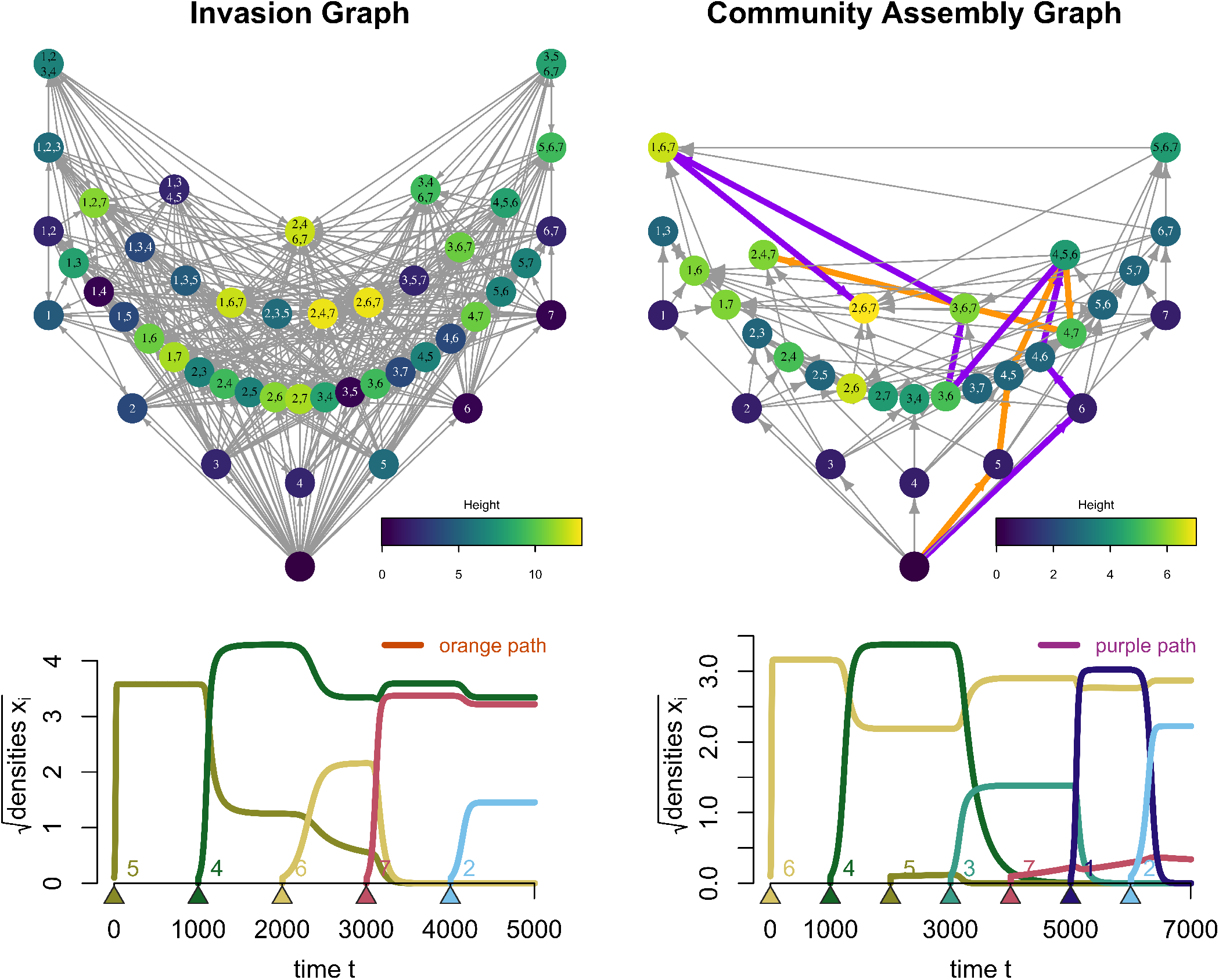
From invasion graphs to assembly dynamics in a 7 species grassland community for an empirically parameterized Lotka-Volterra model by Dormann [2008]. The invasion graph (upper left panel) is acyclic and includes 4 species feasible equilibria. These equilibria correspond to non-permanent and “garden of Eden” communities that are trimmed from the community assembly graph (upper right panel). Sample longest assembly paths are highlighted in orange and in purple. The numerically simulated assembly dynamics of these paths, orange and purple respectively, are shown in the lower two panels. Triangles and labels indicate the timing of species invasions. Parameterization: Species 1 through 7 are *Agrostis capillaris, Festuca rubra, Hydrocotyle heteromeria, Holcus lanatus, Prunella vulgaris, Ranunculus repens*, and *Trifolium repens. A* and *b* matrices are in Supplement C.

There are two three species end-states ({2, 4, 7} and {2, 6, 7}) in the community assembly graph. Despite having the same species richness, these communities are assigned different topographic heights as the longest path length for the community {2, 4, 7} is 5 (orange path in the upper right panel and the lower left simulations of Fig. 3 ) while the longest path length for the community {2, 6, 7} is 7 (purple path and lower right simulations). These maximal paths involve a mixture of invasions that increase species richness and invasions that trigger displacement, reducing richness. As predicted by our mathematical results, both of these communities can be reassembled by introducing each of their species exactly once. For example, 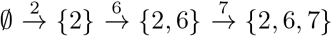 and 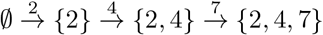.

By our mathematical results, these two end states are alternative stable states: each is locally attracting for the full seven-species dynamics. A community that has reached either state cannot be invaded by the remaining species at low density. Which state is reached is determined by the order of arrival, a priority effect that can be read directly from the graph. For example, starting from the community {2, 7}, the arrival of species 4 leads to {2, 4, 7} and the subsequent arrival of species 6 fails, whereas the arrival of species 6 first leads to {2, 6, 7} and the subsequent arrival of species 4 fails. Permanence requires that every subcommunity be invadable by some missing species, a condition that extends the mutual invasibility criterion of two-species coexistence theory to multispecies communities [Hofbauer and Sigmund, 1998, Schreiber, 2025]. Since no missing species can invade either end state, the seven species cannot coexist in the sense of permanence. Moreover, as the seven-species community does not support a feasible equilibrium and these two end states are the only communities that cannot be invaded by any missing species, every trajectory with all seven species present converges to one of them. The greater height of {2, 6, 7} reflects the existence of longer assembly routes to this end state, not a larger basin of attraction; the community assembly graph describes basins only near the extinction boundary, where sequential low-density invasions occur.

To explore the impacts of the strength of interspecific competition on community assembly, we multiplied all interspecific coefficients *a*_*ij*_ (*i*≠ *j*) by a positive scalar *F*, where *F* = 1 recovers the original parameterization, *F <* 1 weakens interspecific competition and *F >* 1 strengthens it. Across all values of *F* considered, the model satisfies the bounded dynamics criterion, the invasion graph is acyclic, every vertex corresponds to a permanent community, and all edges in the community assembly graph are dynamically realized – there are no false positives.

Varying *F* reveals a hump-shaped relationship between competitive intensity and the complexity of the assembly topography. When interspecific competition is weak (*F* ≪ 1), every invasion succeeds and increases species richness, so all species subsets are permanent communities and maximal path lengths simply equal species richness. The assembly graph has the maximal number of vertices and edges, but the outcome of every invasion is predictable. When interspecific competition is strong (*F* ≫ 1), few invasions succeed, few subsets can coexist, and displacements are rare; assembly is again relatively simple and predictable, but now because most of the graph has been pruned away. At intermediate competitive strengths, the assembly topography is more complex, with displacement events common and outcomes less predictable. Across this gradient, the numbers of vertices and edges of the community assembly graph decline nearly in proportion, whereas invasion success declines more slowly (Figure 4, lower left). The graph shrinks mainly because fewer communities are permanent rather than because invasions from the remaining communities fail.

**Figure 4.**
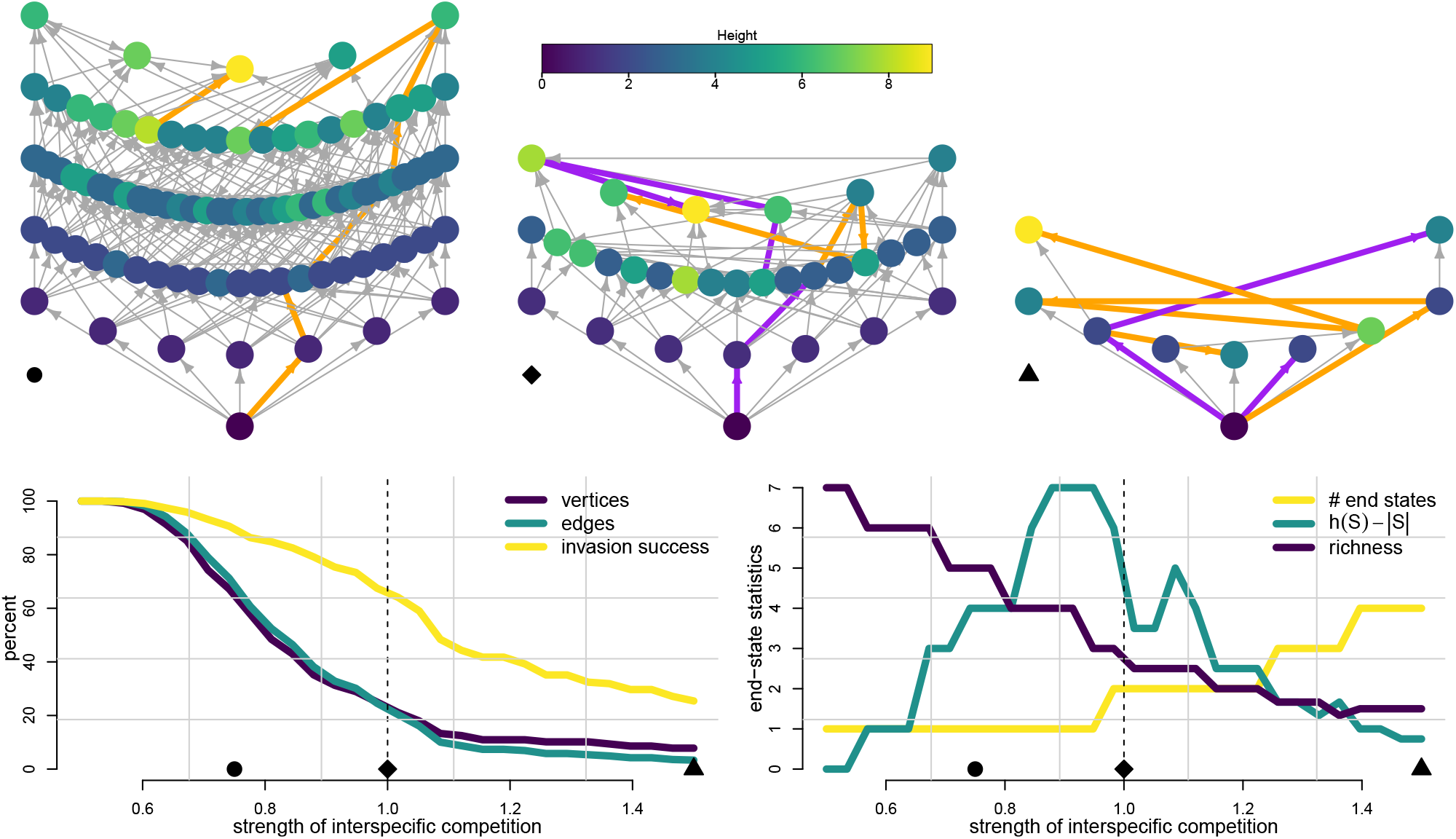
Shifts in the topography of community assembly due to increasing the strength of interspecific competition. *Top:* Community assembly graphs at *F* = 0.75 (circle), *F* = 1 (diamond; the empirical parameterization, shown with labeled vertices in Figure 3), and *F* = 1.5 (triangle). Vertices are arranged vertically by species richness and colored by topographic height; orange and purple edges highlight a longest assembly path to each end state. At *F* = 0.5 (not shown), every subset of species is a permanent community and a vertex of the community assembly graph. *Bottom left:* Numbers of vertices and edges of the community assembly graph as percentages of the 2^7^ subsets of species and of the 7 · 2^6^ possible single-species transitions, respectively, and invasion success, the percentage of missing species with positive invasion growth rates across all feasible communities. *Bottom right:* Number of end states and the mean species richness and mean excess height *h*(*S*) − |*S*| of end states. Dashed lines mark the empirical parameterization *F* = 1; symbols on the horizontal axes mark the values of *F* shown in the top row. Parameters: Baseline (*F* = 1) matrices *A, b* are the same as in Figure 3. For *F*≠ 1, interspecific coefficients *a*_*ij*_ with *i*≠ *j* are multiplied by *F* .

### Facilitative feedbacks in a European pasture

To illustrate how the community assembly graph responds to the strength of interspecific facilitation, we turn to a grassland competition model. Geijzendorffer et al. [2011] estimated seasonal Lotka–Volterra competition coefficients for six grassland species in a managed Irish pasture using empirically derived growth and competition parameters from spring, summer and autumn. We focus on the spring parameterization, which includes six species: *Agrostis stolonifera, Lolium perenne, Phleum pratense, Plantago lanceolata, Trifolium pratense*, and *Trifolium repens*. A notable feature of the spring parameter estimates is that *A. stolonifera* and *P. lanceolata* reciprocally facilitate one another’s growth. Consequently, unlike a purely competitive interaction matrix, the *B*-matrix condition is not immediately satisfied for this model.

Applying the computational pipeline to this model verifies that the *B*-matrix condition holds. The invasion graph (upper left in Figure 5) is acyclic and all vertices correspond to permanent communities. Consequently, our mathematical results imply that only directed edges are trimmed, not vertices, to create the community assembly graph (upper right in Figure 5). This graph includes two complementary 5 species communities: one community {1, 2, 3, 5, 6} has only species 4 missing, and the other community {2, 3, 4, 5, 6} has only species 1 missing. When missing species invade either of these communities, the species 1 and 4, which facilitate one another, exclude all other species. Hence, {1, 4} is the only end state for community assembly—a striking outcome given that five of the six species coexist whenever one member of this pair is absent. The height of this community is 9; the longest assembly paths consist of 9 successful invasions of species. Two of these paths are highlighted in the community assembly graph (orange and purple paths). The pipeline also verifies that all directed edges are dynamically realized. Consequently, consistent with our mathematical results, the numerical simulations of the community assembly recapitulate the changes in community composition predicted by the community assembly graph (lower panels in Fig. 5).

**Figure 5.**
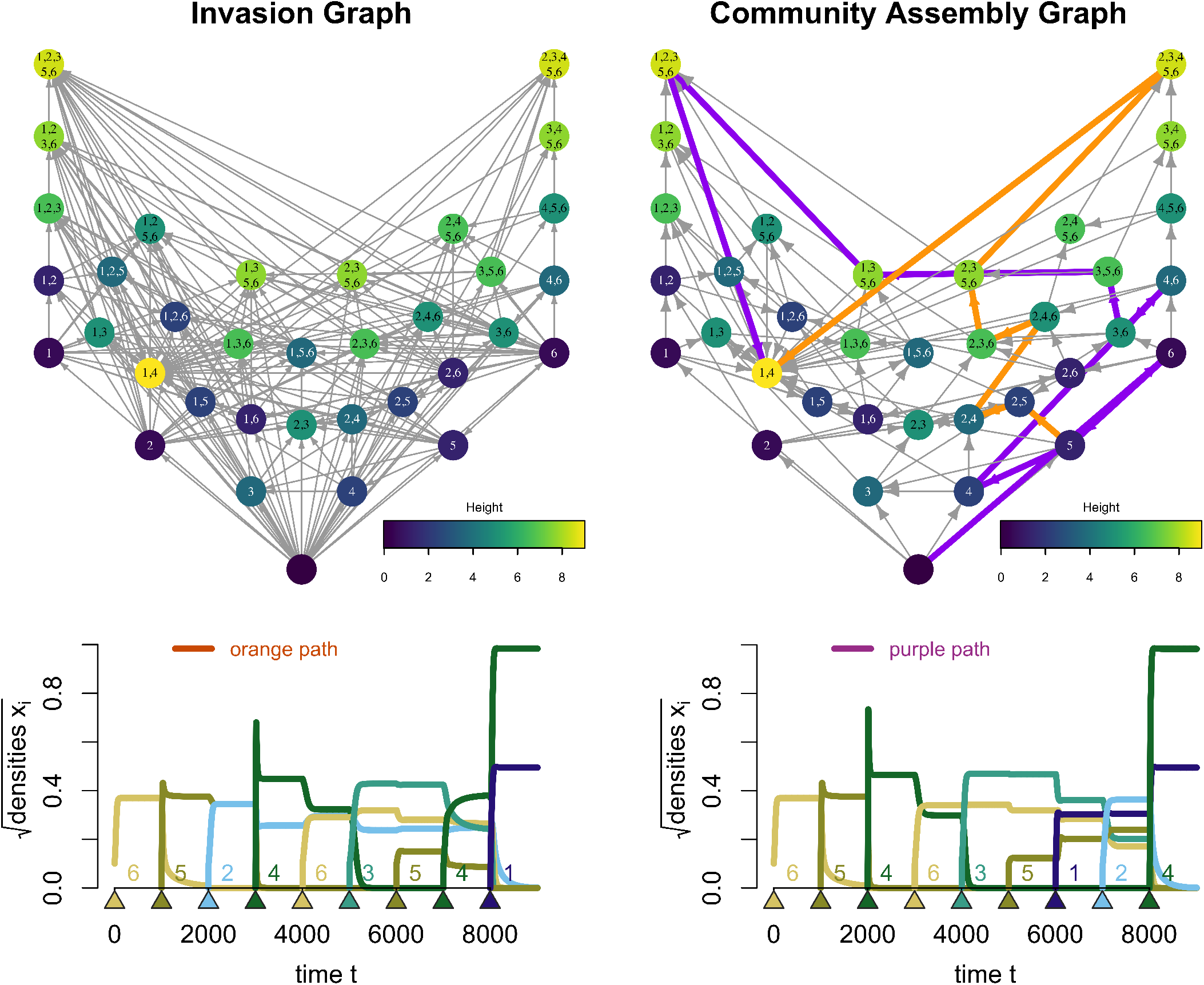
From invasion graphs to assembly dynamics in a 6 species grassland community for an empirically parameterized Lotka-Volterra model by Geijzendorffer et al. [2011]. The invasion graph (upper left panel) is acyclic and all vertices are permanent. Consequently, no vertices are removed for the community assembly graph (upper right panel). Two of the longest assembly paths to the unique end-state are highlighted in orange and in purple. The numerically simulated assembly dynamics of these paths, orange and purple respectively, are shown in the lower two panels. Triangles and labels indicate the timing of species invasions. Parameterization: Species 1 through 6 are *Agrostis stolonifera, Lolium perenne, Phleum pratense, Plantago lanceolata, Trifolium pratense*, and *Trifolium repens. A* and *b* matrices are in supplement C.

To explore how the strength of this mutualism between species 1 and 4 shapes community assembly, we introduce a single parameter *F* ≥ 0 that scales both facilitation coefficients, with *F* = 1 corresponding to empirically estimated values and *F* = 0 for the case where *A. stolonifera* and *P. lanceolata* do not interact. The model satisfies the *B*-matrix condition and, thereby, exhibits bounded dynamics only for *F* ≲ 1.38. This threshold comes from the mutualistic pair alone: the principal submatrix for species 1 and 4 violates the *B*-matrix condition exactly when *F* ^2^*a*_14_*a*_41_ ≥ *a*_11_*a*_44_, that is, when the product of the facilitation coefficients exceeds the product of the self-limitation coefficients. For *F >* 1.38, the positive feedback between the two mutualists overwhelms their self-regulation and their densities grow without bound. For this reason, we restricted *F* to below ≈ 1.38. Across this range of *F* values, the invasion graphs are acyclic, the vertices always correspond to permanent communities, and all edges in the community assembly graph are dynamically realized.

Facilitation strength has little effect on the numbers of vertices and edges of the community assembly graph or on invasion success (Figure 6, lower left). However, it substantially reshapes the end of assembly (Figure 6, lower right). For all values of *F*, assembly terminates in a unique end state, but the richness of this end state declines from five species when the mutualists do not interact (*F* = 0) to the two mutualists alone for *F* ≳ 0.95. Throughout this range, the excess height *h*(*S*) − |*S*| of the end state remains between 5 and 7, so every longest assembly path to this end state involves many species losses.

**Figure 6.**
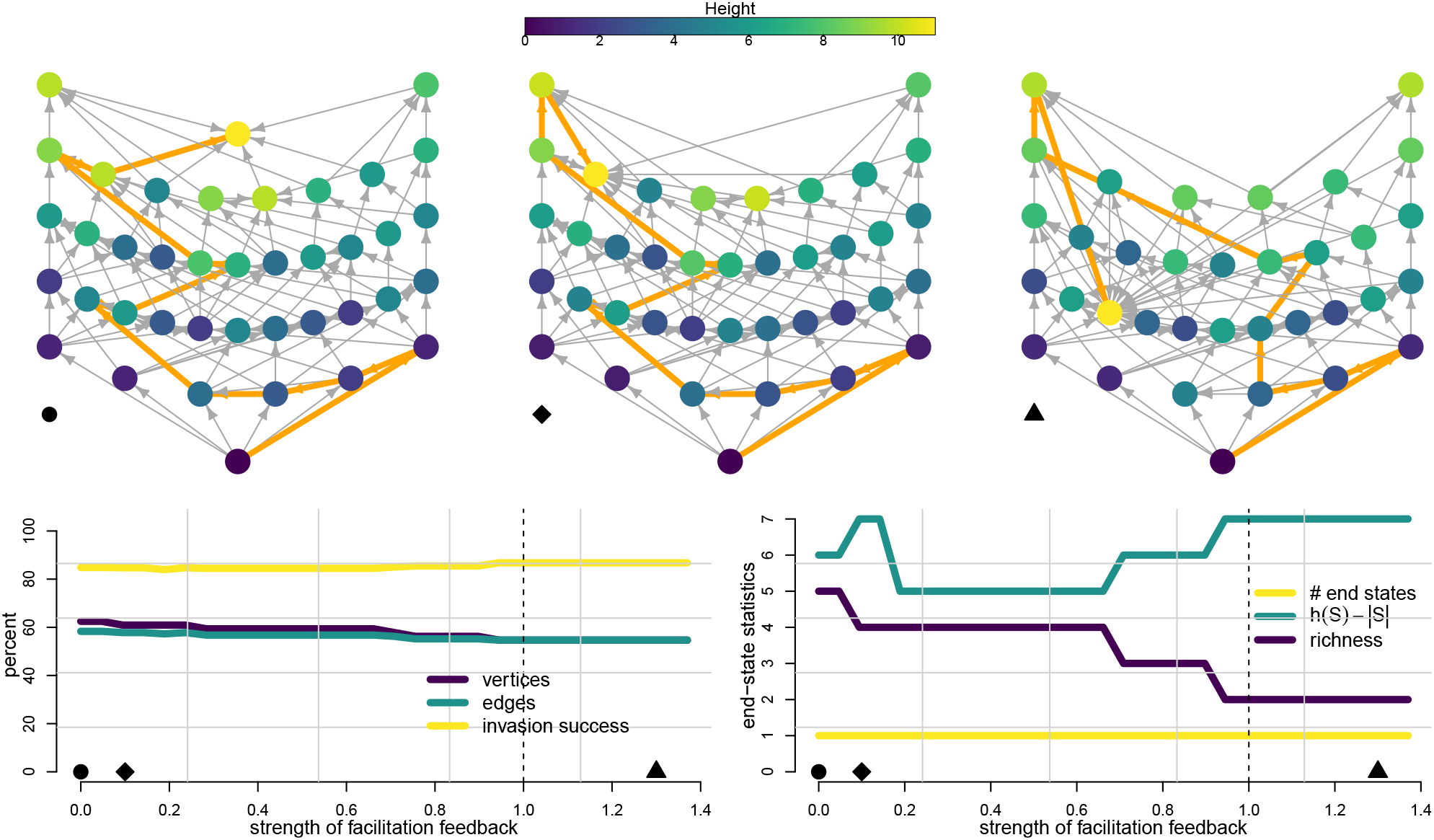
Shifts in the topography of community assembly due to increasing strength of a positive feedback between two mutualists. *Top:* Community assembly graphs at *F* = 0 (circle; no interaction between the mutualists), *F* = 0.1 (diamond), and *F* = 1.3 (triangle); the empirical parameterization *F* = 1 is shown with labeled vertices in Figure 5. Vertices are arranged vertically by species richness and colored by topographic height; orange edges highlight a longest assembly path to the unique end state. *Bottom left:* Numbers of vertices and edges of the community assembly graph as percentages of the 2^6^ subsets of species and of the 6 · 2^5^ possible single-species transitions, respectively, and invasion success, the percentage of missing species with positive invasion growth rates across all feasible communities. *Bottom right:* Number of end states and the species richness and excess height *h*(*S*) − |*S*| of the end state. Dashed lines mark the empirical parameterization *F* = 1; symbols on the horizontal axes mark the values of *F* shown in the top row. Values *F* ≳ 1.38 are excluded because the *B*-matrix condition fails. Parameters: Base line (*F* = 1) matrices *A, b* are the same as in Figure 5. For *F*≠ 1, facilitation coefficients *a*_14_, *a*_41_ are multiped by *F* .

### Cycling in Ant Communities

To illustrate how our methods apply to communities that exhibit assembly cycles, we consider a model motivated by the ant community of a coffee agroecosystem in Puerto Rico studied by Vandermeer and Perfecto [2023]. This system is centered on an intransitive competitive cycle among three invasive ant species—the red imported fire ant *Solenopsis invicta*, the little fire ant *Wasmannia auropunctata*, and the flower ant *Monomorium floricola*—in which *S. invicta* displaces *W. auropunctata, W. auropunctata* displaces *M. floricola*, and *M. floricola* in turn displaces *S. invicta* through direct chemical interference. Field censuses spanning four years document the cycling of these three dominants across coffee bushes, consistent with the rock–paper–scissors dynamics predicted by theory [May and Leonard, 1975]. Connected to this intransitive core is a set of subordinate species presumed to compete transitively, whose persistence is maintained through the competitive opportunities created by the recurrent cycling of the dominant triad. Vandermeer and Perfecto [2023] formalized this structure with a Lotka–Volterra model in which all intrinsic growth rates and intraspecific competition coefficients are set to one (*b*_*i*_ = 1, *a*_*ii*_ = −1), the intransitive triad (*N*_1_, *N*_2_, *N*_3_) competes with a common interspecific coefficient −*a*, and each subordinate species experiences competitive pressure from its anchoring triad member with coefficient −*a* and from other subordinates through a hierarchical coefficient *h* ≥ *a*. This model provides a natural test case for our invasion graph methodology in the presence of cycles in the invasion graph, complementing the empirically parameterized models considered elsewhere in this section.

The invasion graph for this model contains directed cycles, reflecting the intransitive dynamics of the dominant triad. For communities lacking at least one member of the triad, the invasion graph restricted to the community is acyclic, and Theorem 3 establishes permanence. For communities containing the full triad, the restricted invasion graph is cyclic and the invasion graph criterion is only necessary. However, each of these communities satisfy the permanence criterion of Hofbauer [1981] (Supplement B). Hence every vertex of the invasion graph is permanent. None the less, communities containing the full triad cannot be assembled by the sequential single-species invasions that define the community assembly graph. Consequently, they are removed in Step 4 of the pipeline. Every vertex of the resulting community assembly graph is permanent by the acyclic sufficient condition, and all of its edges are dynamically realized.

The resulting community assembly graph exhibits a complex network of intertwined cycles, two of which are highlighted in Figure 7 along with simulations of the corresponding Lotka-Volterra dynamics. Both simulations trace heteroclinic cycles that involve the dominant triad. Each transition along these cycles is a dynamically realized single-species invasion and corresponds to a connecting orbit between successive communities. In the simulations, species that fall below the extinction threshold are removed and later reintroduced at low density, which allows the assembly dynamics to traverse these cycles repeatedly. These assembly cycles capture, at the level of community composition, the same rock-paper-scissors turnover documented in the field.

**Figure 7.**
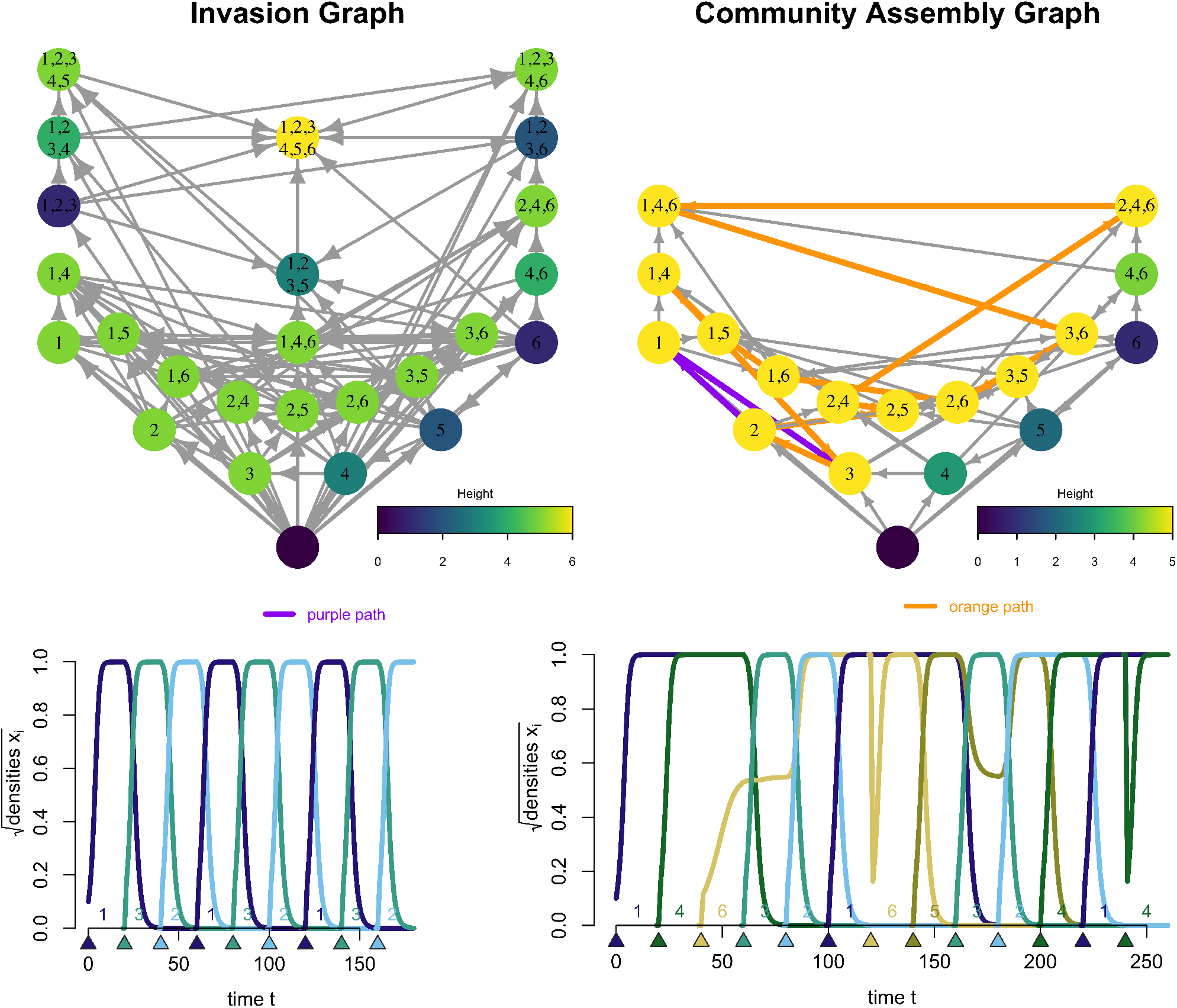
From invasion graphs to assembly dynamics in a 6 species ant community of a coffee agroecosystem for an empirically inspired model by Vandermeer and Perfecto [2023]. The invasion graph (upper left panel) is cyclic and every vertex is permanent: for vertices lacking at least one member of the intransitive triad, permanence follows from the acyclic sufficient condition of Theorem 3; for vertices containing the full triad, permanence is verified using the criterion of Hofbauer [1981] (Supplement B). The latter communities cannot be assembled by single-species introductions and are removed from the community assembly graph (upper right panel). Two assembly cycles are highlighted in purple and orange, and the numerically simulated assembly dynamics along them are shown in the lower left and lower right panels, respectively. Triangles and labels indicate the timing of species invasions. Parameterization: *A* and *b* matrices as described in main text.

## Discussion

Community assembly graphs provide a convenient summary of which subsets of species can coexist and how single-species invasions drive transitions between these subcommunities. When built from dynamical models, they connect the assembly process to the underlying ecological mechanisms. However, realizing this promise requires constructing the graph in a way that is computationally tractable and faithful to the underlying dynamical model. Here, we developed a pipeline that achieves this. In this discussion, we consider the methodological ground it gains over existing approaches, the ecological meaning it gives the assembly landscape metaphor, the insights our empirical case studies provide, and the challenges and opportunities that remain.

### Methodological advances

Our pipeline offers five advances: it verifies that community dynamics are bounded, identifies which subsets of species coexist in the sense of permanence using invasion graphs, identifies the edges of the invasion graph corresponding to single-species invasions, identifies the vertices of the invasion graph corresponding to communities that are assembled by single-species introductions, and constructs community assembly graphs that are mathematically guaranteed to reproduce the dynamics of assembly under rare single-species invasions. Together, these make the pipeline a tool an empiricist can apply directly to a Lotka–Volterra parameterization. This extends the reach of earlier dynamical approaches to CAG construction. The methods of Serván and Allesina [2021] and Godoy et al. [2024] require global stability or VL-stable interaction matrices, assumptions that exclude alternative stable states and non-equilibrium dynamics. Because the time-averaging property makes invasion growth rates independent of the resident attractor, our pipeline accommodates both without ever computing the attractors themselves. Both of these dynamics arise in empirically parameterized models: our grassland example exhibits alternative states [Roxburgh, 1994, Roxburgh and Wilson, 2000, Dormann, 2008], and our ant community example exhibits non-equilibrium dynamics [Vandermeer and Perfecto, 2023]. Our dynamical-realization check is automatically satisfied under the assumption of a globally stable equilibrium. Therefore, since global stability is a special case of permanence, the result of Serván and Allesina [2021], Godoy et al. [2024] are a special case of our framework.

Earlier work [Law and Blackford, 1992, Law and Morton, 1996] also equated coexistence with permanence. However, their algebraic verification of permanence is more restrictive and computationally more involved. Law and Morton [1996] used the criterion of Hofbauer [1981], which requires the existence of positive weights *p*_*i*_ with 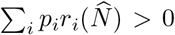 at every feasible equilibrium 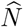 with at least one missing species. Jansen [1987] showed that this condition can be verified by solving a linear programming problem, a computational step that the invasion graph criterion avoids. The two criteria differ in how reliably they identify permanence. For acyclic invasion graphs, the invasion graph criterion is sharp: it certifies permanence if and only if the community is permanent. The Hofbauer criterion is not. It can fail to certify genuinely permanent communities even in this acyclic case, e.g. predators that mediate permanence of founder-controlled prey [Schreiber, 1997, Hening et al., 2022]. For cyclic invasion graphs, neither criterion fully characterizes permanence, but the two can be combined to expand the set of communities whose permanence can be resolved [Schreiber, 2000, Spaak and Schreiber, 2023]. Our ant example illustrates this complementarity: the invasion graph criterion certifies permanence for every community lacking at least one dominant, and the criterion of Hofbauer [1981] certifies the remaining communities containing the intransitive triad (Supplement B). A systematic integration of the two criteria for general cyclic invasion graphs remains an open problem.

Verifying that community dynamics remain bounded is a step that none of the earlier methods checks explicitly, yet it must hold for the invasion graph to be meaningful. For interaction matrices built from competition and predator–prey interactions, boundedness is typically automatic, e.g. the hierarchical condition in Hofbauer and Sigmund [1998], which may explain why it has received little attention. Once positive interactions enter, however, it can fail, and in less obvious ways than one might expect. It is well known that mutualistic models can exhibit unbounded growth [May and McLean, 2007] as in our European pasture example [Geijzendorffer et al., 2011]. However, blow-up can also arise from positive feedback loops involving more than two species, even when every pair of species has bounded dynamics. For example, in the Lotka–Volterra dataset of Spaak et al. [2021], a 9 species model based on Caputa [1948] contains a three-species subcommunity whose dynamics are unbounded even though every one- and two-species subset is bounded (Supplement D, Fig. D0). Our inductive *B*-matrix test detects exactly these cases, and identifying them is a prerequisite for applying the framework to communities with facilitation or other positive interaction. This is an important extension given the increasing recognition that facilitation is common in what were once thought to be purely competitive communities [Suding et al., 2024, James et al., 2025].

### The Assembly Landscape

Evolution as climbing a fitness landscape, though an imperfect metaphor, has been a powerful organizing idea in the development of evolutionary theory. Several authors have sought a similar metaphor for community assembly [Luh and Pimm, 1993, George and Korolev, 2023]. George and Korolev [2023], for example, draws an explicit analogy in which community function plays the role of fitness and community composition the role of the genome. Luh and Pimm [1993] went further and assigned each community state a “height, a fitness value, if you like,” with assembly always moving from a lower state to a higher neighbor. They were, however, openly skeptical of their own construction, worrying that it made the community behave like “some superorganism ever driven to greater perfection,” and concluding that they had “no idea if we should model community assembly graphs by landscapes, let alone what kind of landscape is the correct one.” Despite these legitimate concerns, we show that there is an ecologically meaningful topography for both acyclic and cyclic community assembly graphs. For an acyclic graph, the height of a vertex is the length of the longest assembly path of single-species invasions leading to that community. For a cyclic graph, all communities along a cycle are assigned a common height, equal to the longest assembly path reaching any community in the cycle. Within a strongly connected component there is no consistent sense in which one member is higher than another, which is precisely why the component is collapsed to a single elevation. One can picture these components as interlocking ridges at equal elevation, locally higher than the off-ridge communities that feed into them.

The ecological content of this topography lies in the way height relates to species richness. Because height is bounded below by richness, the two coincide exactly when every invasion along the longest path adds a species, and none triggers a displacement. This displacement-free limit is the Clementsian “superorganism” [Clements, 1916]: assembly proceeds by strict accumulation, and height simply counts the species present. When height exceeds richness, on the contrary, the longest assembly paths contain invasions that displace one or more residents, and the excess *h*(*S*) − |*S*| counts the residents lost along such a path, an invasion that displaces two residents counting twice. A community organized by a competitive hierarchy illustrates the point: dominants sit higher in the landscape than the subordinates they displace, so height tracks position in the hierarchy rather than richness. Height is an extremal statistic, a property of the longest path rather than of a typical one. Under the conditions of our minimal assembly result, however, every community *S* in the graph can also be reached by a path of length |*S*|, so *h*(*S*) − |*S*| is the full range of assembly path lengths to *S*. Unlike the mean or other moments of path length, the range does not depend on a probability distribution over introduction sequences, so it summarizes the historical contingency of assembly in a way that is determined by the interaction parameters alone. This topography provides one resolution to Pimm’s open question of “what the abstraction ‘akin to height’ is, I do not know” [Pimm, 1991, p. 251].

### New empirical insights

Applying the CAG pipeline to the empirically parameterized and empirically inspired Lotka–Volterra models highlights several points. First, in the grassland model of Dormann [2008], many of the feasible equilibria of the invasion graph are pruned in constructing the community assembly graph. Because the invasion graph is acyclic, our results guarantee that all of these pruned vertices correspond to non-permanent communities. Equating coexistence with the mere existence of a feasible equilibrium [Saavedra et al., 2017] could therefore lead to misleading inferences about community assembly, and this underscores why the full invasion graph must be computed as the first step in producing the community assembly graph. The same example exhibits two alternative stable states reached through a priority effect between two species, and the graph identifies both the states and the assembly sequences that select between them, without any assumption of global stability. Moreover, this example highlights how the assembly topography changes as interspecific competition is varied, from strong negative frequency dependence (weak interspecific competition) to strong positive frequency dependence (strong interspecific competition). At either extreme, the topography reduces to species richness: every invasion succeeds when competition is weak, whereas essentially no invasion succeeds once a single species is established when competition is strong. Between these extremes, the topography becomes markedly more complex. Second, the pasture model of Geijzendorffer et al. [2011] shows how positive feedbacks can produce unbounded growth, and also yields a “1066 effect” [Lockwood et al., 1997]: a pair of mutualists that together displace all other species, yet coexist with the rest of the community whenever either member of the pair is absent. Finally, the ant community model of Vandermeer and Perfecto [2023] provides an empirically inspired instance of the “Humpty Dumpty” effect [Pimm, 1991], in which subcommunities that are permanent cannot be assembled by any sequence of single-species introductions and must instead be introduced simultaneously. Law and Morton [1996] termed such communities “fragile”, and here they arise directly from the intransitive competition of the dominant triad.

### Future opportunities and challenges

The pipeline opens the door to studying assembly across a much larger set of empirically parameterized Lotka–Volterra models, and with it a number of classical questions. How common are multiple alternative end states, as opposed to a single climax community? To what extent do species interactions constrain the way local richness scales with the size of the regional species pool? And how often do transient priority effects arise, in which the order of species introductions determines the outcome of assembly? Beyond individual systems, the assembly topography invites comparative analysis: whether there are cross-system regularities in landscape ruggedness or in the distribution of community heights, connecting the framework to macroecological questions about the predictability of assembly. Complementing these broad comparisons, modern coexistence theory offers a route to the mechanisms behind assembly in particular systems, clarifying why individual invasions succeed, fail, or trigger displacement [Brennan and Schreiber, 2026].

Several methodological challenges remain. The first challenge concerns directed cycles in the invasion graph. Verifying permanence in their presence is hard in general, but for “simple” cycles [Hofbauer and So, 1994] the invasion-graph and Hofbauer criteria can be combined in the spirit of Schreiber [2000]. Complex cycles, in which cycles are nested within other cycles [Brannath, 1994], raise open mathematical questions that new methods will be needed to resolve. A second, more representational challenge arises when a species can successfully invade a community, yet the invasion leads not to a new community but to a cycle. How such transitions are best depicted in a community assembly graph is an open challenge. A further frontier is extending the framework beyond Lotka-Volterra dynamics. Invasion-graph methods already exist for models with nonlinear per-capita growth rates, impulsive dynamics, temporal forcing, eco-evolutionary feedbacks, and population structure [Schreiber, 2025]. While the conceptual path is clear, the practical obstacle is that invasion growth rates become far harder to compute once the time-averaging property no longer holds. Finally, an important challenge is extending this framework to alternative notions of coexistence, most notably the existence of a positive attractor. This requires going beyond both the invasion-growth-rate criterion underlying permanence and the locally stable equilibrium criterion used elsewhere [Schreiber, 2006]. The choice of coexistence concept is not simply a technicality. Under permanence the “resident strikes back” phenomenon cannot occur, yet Case [1995] showed it appears once coexistence is equated with a locally stable equilibrium. This sensitivity is a concrete reminder that what one means by coexistence shapes what one concludes about assembly as the framework moves beyond permanence.

A practical consideration is how the computational complexity of the pipeline scales with the number of species *k*. Steps 1 and 2 examine every subset of the species pool, computing a determinant and adjoint (Step 1) or solving a linear system (Step 2) of size at most *k* per subset, for a total of *O*(2^*k*^*k*^3^) operations. Step 2 then tests every ordered pair of feasible equilibria for a directed edge, a cost of *O*(*F* ^2^*k*) where *F* is the number of feasible equilibria. When most subsets are feasible, as under weak interspecific competition, *F* approaches 2^*k*^ and this edge test, at *O*(4^*k*^*k*), dominates the subset enumeration. When many subsets are infeasible, as in our grassland example where only 46 of 128 subsets support a feasible equilibrium, the subset enumeration dominates. The remaining steps of the pipeline are graph algorithms whose cost is linear in the number of edges, which is at most *F* (*F* −1)*/*2 and equals 3^*k*^ − 2^*k*^ when every subset is feasible and every missing species can invade. Memory requirements are *O*(*Fk*) for the equilibria and invasion growth rates and at most *O*(*F* ^2^) for the edges. In our R implementation (R version 4.6.0 on a MacBook Pro with an Apple M4 chip), the seven-species grassland model runs in 0.005 seconds. For random competitive communities with *b*_*i*_ = 1, *a*_*ii*_ = −1, and *a*_*ij*_ = −|*z*_*ij*_| for independent standard normal *z*_*ij*_, median run times across ten communities of each size were 0.05 seconds (range 0.02–0.09) for *k* = 10, 0.68 seconds (range 0.50–0.99) for *k* = 15, and 20.3 seconds (range 14.9–45.0) for *k* = 20, with medians of 117, 827, and 7,044 feasible equilibria, respectively. For the empirically parameterized Lotka-Volterra models that motivate this work, which rarely exceed 20 species, exhaustive construction is therefore feasible. For larger pools, the community assembly graph could be constructed forward from the empty community, computing only communities reachable by single-species invasions, in which case the cost is governed by the richness of the largest reachable community rather than by *k*. Developing this type of inductive approach is a natural next step.

### Conclusion

Community assembly is not only a question of which communities are possible, but of what paths lead there and how the topography of those paths structures historical contingency. The methods introduced here make this topography quantifiable in a mathematically justified manner.

## Supplement A. Mathematical Details for the Community Assembly Graph

This Supplement provides the mathematical and computational details underlying the community assembly graph (CAG) pipeline, the mathematical properties of CAGs, and the numerical simulations of assembly in the Lotka-Volterra (LV) models. Throughout, we consider LV models of the form

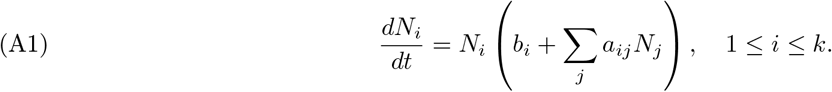

This supplement is organized as follows. We begin with the CAG pipeline, proceeding in logical order: verifying that the model has uniformly bounded dynamics, computing the invasion graph and using it to verify permanence of candidate communities, and establishing the mathematical justification for identifying single-species transitions and confirming that directed edges in the CAG are dynamically realized. We then prove three properties of CAGs stated in the main text: that a successful invader always persists in the resulting community, that every community in the CAG can be assembled via a path whose length equals its species richness, and that end states corresponding to permanent communities are dynamical attractors. This supplement concludes with the numerical algorithm for simulating assembly sequences and a theorem rigorously connecting those simulations to paths in the CAG.

## The Community Assembly Pipeline

### Verifying Boundedness

A Lotka-Volterra model is *uniformly bounded* if there exists *M >* 0 such that lim sup_*t*→∞_ max_*i*_ *N*_*i*_(*t*) ≤ *M* whenever min_*i*_ *N*_*i*_(0) ≥ 0. Uniform boundedness ensures that the dynamics remain ecologically meaningful and is a prerequisite for the invasion graph to be well-defined. An interaction matrix *A* is a *B-matrix* if the associated Lotka-Volterra model is uniformly bounded for all choices of the intrinsic growth rate vector *b*. Hofbauer and Sigmund [1998] established the following equivalent conditions for *A* to be a *B*-matrix.

#### Theorem 1

(Hofbauer and Sigmund [1998]). *The following are equivalent:*

i. *A is a B-matrix*.
ii. *Whenever N*_*i*_ *∑* _*j*_ *a*_*i j*_ *N*_*j*_ = *λN*_*i*_ *for all* 1 ≤ *i* ≤ *k and some non-zero N* ≥ 0, *then λ <* 0.
iii. *For all non-zero N* ≥ 0, *∑* _*j*_ *a*_*ij*_*N*_*j*_ *<* 0 *for some i with N*_*i*_ *>* 0.

To verify the *B*-matrix condition algorithmically, Hofbauer and Sigmund [1998] provide an inductive criterion based on principal submatrices. Recall that the **adjoint** of a matrix *A*, denoted adj(*A*), is the matrix whose (*i, j*)-entry is (−1)^*i*+*j*^ times the determinant of the (*n* − 1) × (*n* − 1) submatrix obtained by removing the *j*-th row and *i*-th column of *A*. For an invertible matrix, *A*^−1^ = adj(*A*)*/* det(*A*), so condition (b) below is equivalent to all entries of *A*^−1^ being positive.

#### Theorem 2

(Hofbauer and Sigmund [1998]). *If all* (*n* − 1) × (*n* − 1) *principal submatrices of A are B-matrices, then the following are equivalent:*

a. *A is not a B-matrix*.
b. det(−*A*) ≤ 0 *and* adj(−*A*) *>* 0.

Algorithm 1 applies Theorem 2 inductively to verify whether *A* is a *B*-matrix, checking principal submatrices of increasing size and terminating as soon as a violation is detected.

### Computing the Invasion Graph and Verifying Permanence

Building on the verified boundedness of the model, we now compute the invasion graph and use it to determine which candidate communities are permanent. We assume throughout this subsection that the Lotka-Volterra model is uniformly bounded and has a finite number of equilibria, i.e., all principal submatrices of *A* are invertible. Under these conditions, Hofbauer and Schreiber [2022] introduced the invasion graph *G* associated with the model. The vertices of *G* correspond to

#### Algorithm 1.

*B*-matrix verification for a *k* × *k* interaction matrix *A*

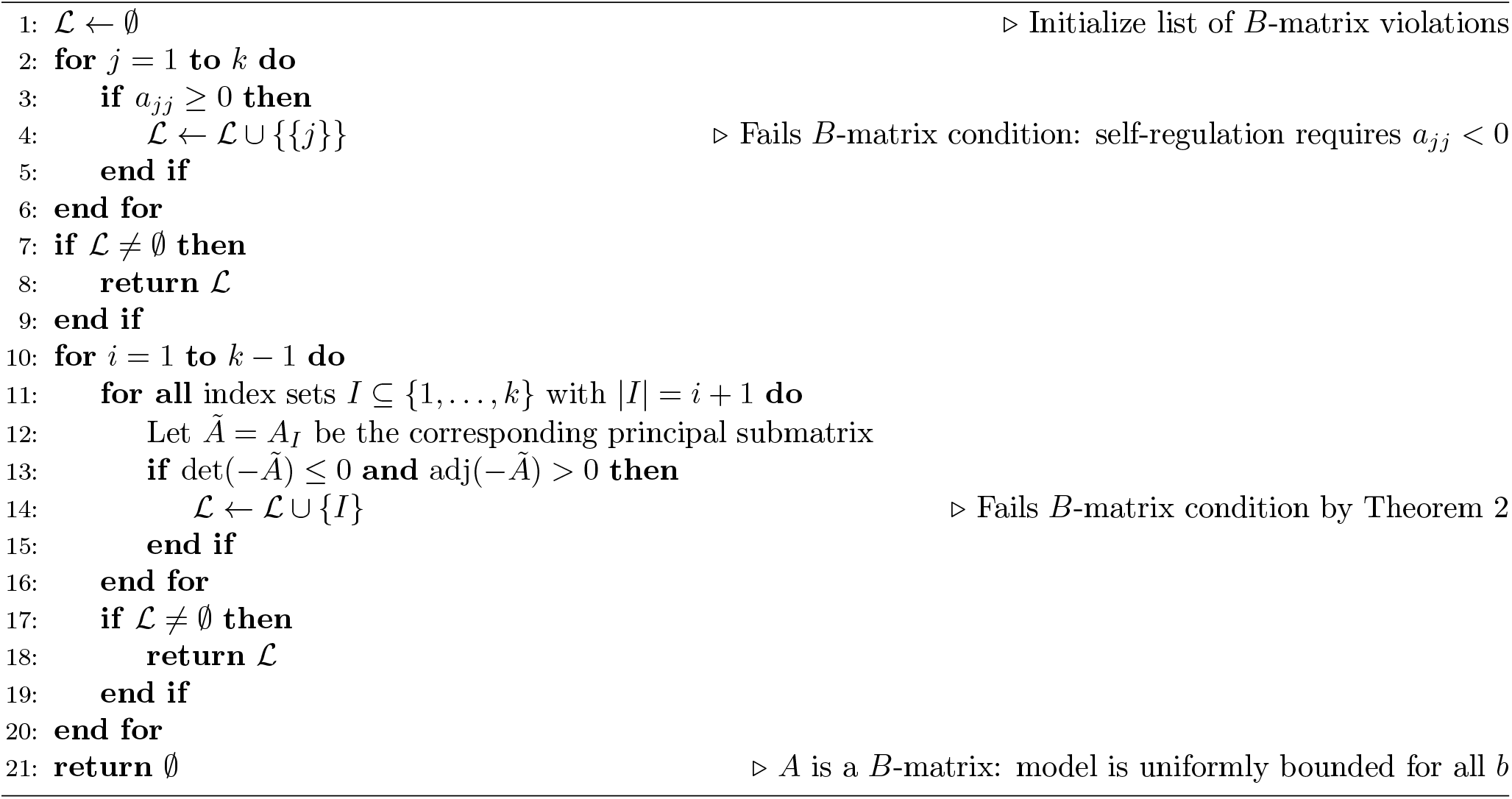

subsets *S* ⊆ {1, 2, …, *k*} for which there exists an equilibrium *x*^*^(*S*) with 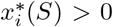 if and only if *i* ∈ *S*. We denote the collection of all such subsets by *S*. The per-capita growth rate of species *i* at equilibrium *x*^*^(*S*) is 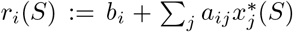, with *r*_*i*_(*S*) *>* 0 indicating that species *i* can invade community *S* and *r*_*i*_(*S*) *<* 0 indicating that it cannot. There is a directed edge *S* → *T* in *G* if *r*_*i*_(*S*) *>* 0 for all *i* ∈ *T* \ *S* and *r*_*i*_(*T* ) *<* 0 for all *i* ∈ *S* \ *T* .

The invasion graph serves two purposes in the CAG pipeline. First, and most importantly, for our purposes, it can be used to verify the permanence of candidate communities. A community *S* is *permanent* if all species in *S* persist indefinitely regardless of initial conditions, i.e. there exists *m >* 0 such that lim inf_*t*→∞_ *N*_*i*_(*t*) ≥ *m* for all *i* ∈ *S* whenever *N*_*i*_(0) *>* 0 for all *i* ∈ *S*. Hofbauer and Schreiber [2022] proved the following theorem, which can be applied to any subset of species by restricting the model to that subset.

#### Theorem 3

(Hofbauer and Schreiber [2022]). *Let* S *be the collection of proper subsets S* ⊊ {1, …, *k*} *with a corresponding equilibrium x*^*^(*S*).

#### Necessary condition for permanence

*If the community* {1, …, *k*} *is permanent, then* max_*i*_ *r*_*i*_(*S*) *>* 0 *for all S* ∈ S. *That is, at least one missing species can invade every proper subcommunity*.

#### Sufficient condition for permanence

*If G is acyclic and* max_*i*_ *r*_*i*_(*S*) *>* 0 *for all S* ∈ S, *then the community* {1, …, *k*} *is permanent. That is, the necessary condition is also sufficient when the invasion graph is acyclic*.

Second, the invasion graph identifies the connecting orbits between communities. If there exists a solution *N* (*t*) such that

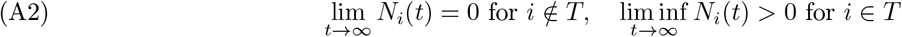

and

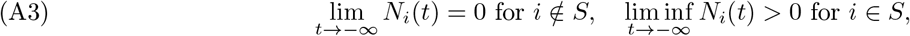

then *S* → *T* is a directed edge in *G*. Such solutions are called *connecting orbits* in the dynamical systems literature and correspond to transitions between communities driven by invasion dynamics. Algorithm 2 describes the computation of *G*.

##### Algorithm 2.

Invasion graph computation for a Lotka-Volterra model

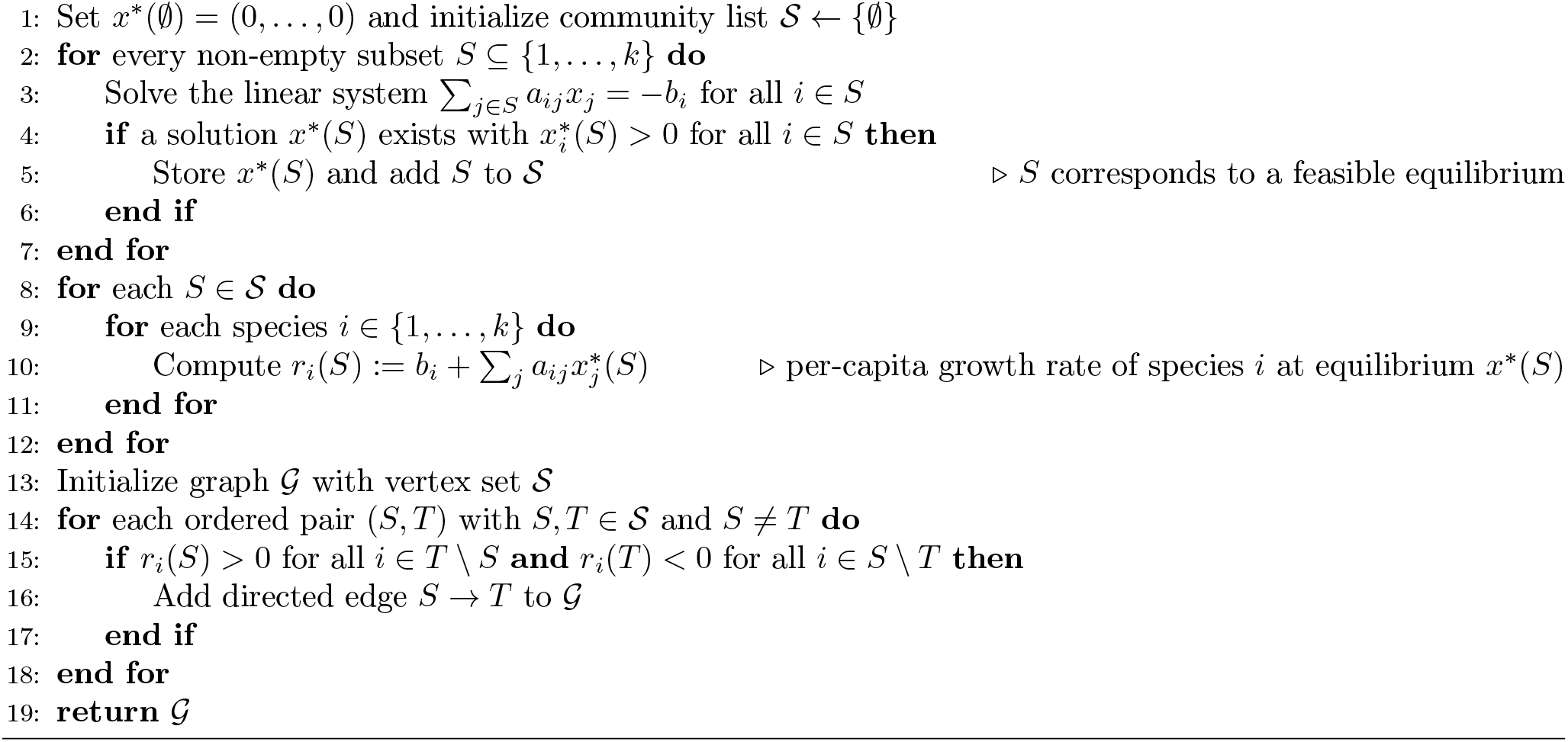

### Pruning and Verifying Dynamical Realization

The CAG is constructed from the invasion graph G by retaining only edges that correspond to single-species invasions and verifying that each such edge is dynamically realized. This subsection establishes two results that justify this procedure. Proposition A.1 shows that a positive invasion growth rate together with acyclicity of the restricted invasion graph guarantees the existence of a single-species transition out of *S*. Proposition A.2 shows that when the resulting community is the unique saturated community in *S* ∪ {*i*}, the dynamics converges to that community from all initial interior conditions in *S* ∪ {*i*}. We say that a community *T* ⊆ *S* ∪ {*i*} is *saturated* in *S* ∪ {*i*} if *r*_*j*_(*T* ) *<* 0 for all *j* ∈ (*S* ∪ {*i*}) \ *T*, i.e., no missing species from the pool *S* ∪ {*i*} can invade *T* .

#### Proposition A.1.

*If r*_*i*_(*S*) *>* 0 *and the invasion graph* G *restricted to S* ∪ {*i*} *is acyclic, then there exists T* ∈ S *such that* 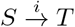.

*Proof*. As the proof relies on results from Hofbauer and Schreiber [2022], we adopt their notation and refer the reader there for definitions. Since *r*_*i*_(*S*) *>* 0, the unstable manifold of the equilibrium *x*^*^(*S*) intersects the set {*N* : *N*_*j*_ *>* 0 iff *j* ∈ *S* ∪ {*i*}}. Acyclicity of the invasion graph restricted to *S* ∪ {*i*} and [Hofbauer and Schreiber, 2022, Lemma 4] imply that there exists *T* ∈ S with *T* ⊆ *S* ∪ {*i*} such that the *ω*-limit set of this unstable manifold lies in {*N* : *N*_*j*_ *>* 0 iff *j* ∈ *T* }. [Hofbauer and Schreiber, 2022, Lemma 2] then implies 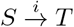 . □

Proposition A.1 does not guarantee that *T* is unique, or that the transition 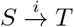 is dynamically realized. However, the following proposition guarantees both properties.

#### Proposition A.2.

*If* 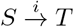, *the invasion graph restricted to S* ∪ {*i*} *is acyclic, and T is the unique saturated community in S* ∪ {*i*}, *then there exists m >* 0 *such that*

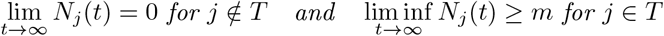

*whenever N*_*j*_(0) *>* 0 *for j* ∈ *S* ∪ {*i*} *and N*_*j*_(0) = 0 *otherwise*.

*Proof*. As the proof relies on results from Hofbauer and Schreiber [2022], we adopt their notation and refer the reader there for definitions. First, suppose *T* = *S* ∪ {*i*}. The invasion graph restricted to *T* is acyclic and every proper subcommunity 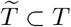 has at least one species 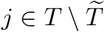 with 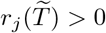. [Hofbauer and Schreiber, 2022, Theorem 1] then implies that *T* is a permanent community, so there exists *m >* 0 such that lim inf_*t*→∞_ *N*_*j*_(*t*) ≥ *m* whenever *N*_*j*_(0) *>* 0 for all *j* ∈ *T* and *N*_*j*_(0) = 0 otherwise.

Now suppose *T* is a proper subset of *S* ∪{*i*}. Since *T* is the unique saturated community in *S* ∪{*i*}, there is no equilibrium in *C*_+_ := {*N* : *N*_*j*_ *>* 0 iff *j* ∈ *S* ∪ {*i*}}. Let *C* := {*N* : *N*_*j*_ = 0 for all *j* ∈*/ S* ∪ {*i*}}. Since the invasion graph restricted to *S* ∪ {*i*} is acyclic, [Hofbauer and Schreiber, 2022, Lemma 4] implies *ω*(*N* (0)) ⊂ *C* \ *C*_+_ for all *N* (0) ∈ *C*_+_. Since *T* is the unique saturated community in *S* ∪ {*i*}, [Hofbauer and Schreiber, 2022, Lemma 4] implies that there exists a compact set *K* ⊂ {*N* : *N*_*j*_ *>* 0 iff *j* ∈ *T* } such that *ω*(*N* (0)) ⊂ *K* for all *N* (0) ∈ *C*_+_. By compactness of *K*, there exists *m >* 0 that satisfies the claimed convergence whenever *N* (0) ∈ *C*_+_. □

## Properties of the Community Assembly Graph

This section establishes three properties of CAGs that are stated in the main text. Proposition A.3 shows that a successful invader always persists in the resulting community, provided that the resident community is permanent. Proposition A.4 shows that every community in the CAG can be assembled through a path whose length equals its species richness, i.e., one introduction per species. Proposition A.5 shows that end states corresponding to permanent communities are dynamical attractors and that a unique end state with an acyclic invasion graph attracts all positive initial conditions.

### Proposition A.3.

*Let* 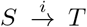 *be a directed edge in the community assembly graph. If S is a permanent community, then i* ∈ *T* .

*Proof*. Assume to the contrary that *i* ∈*/ T* . Then *T* ⊆ *S* and *r*_*j*_(*T* ) *<* 0 for all *j* ∈ (*S* ∪ {*i*}) \ *T* . In particular, *r*_*i*_(*T* ) *<* 0. Since *r*_*i*_(*S*) *>* 0 and *r*_*i*_(*T* ) *<* 0, *T* is a proper subset of *S* and max_*j*∈*S*\*T*_ *r*_*j*_(*T* ) ≤ 0. However, since *S* is a permanent community, Theorem 3 implies max_*j*∈*S*\*T*_ *r*_*j*_(*T* ) *>* 0, a contradiction. □

### Proposition A.4.

*Let S be a permanent community such that the invasion graph restricted to S is acyclic. Then there is a directed path of single-species transitions from* ∅ *to S whose vertices are subcommunities of S, so S is not removed in Step 4. If, in addition, Step 5 prunes no edge between subcommunities of S, then S is in the community assembly graph and, if all the vertices of the community assembly graph are permanent, there is a directed path of length* |*S*| *from* ∅ *to S in the community assembly graph*.

When the invasion graph is cyclic and *S* is permanent, *S* may not be in the community assembly graph. For example, the classical rock-paper-scissor example of May and Leonard [1975] has a cyclic invasion graph and allows for permanence of the three competing species. However, in this case, the three species community cannot be reconstructed through single-species introductions. Pimm [1991] called this outcome the “Humpty Dumpty effect”.

*Proof*. Assume that the invasion graph is acyclic and *S* is permanent. We prove that there exists a directed path of single-species invasions from ∅ to *S* and, consequently, *S* lies in the community assembly graph. Since *S* is permanent, Theorem 3 implies 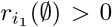 for some *i*_1_ ∈ *S*, and Proposition A.1 implies that there exists *T*_1_ ⊆ *S* such that 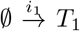 . If *T*_1_ = *S* we are done. Otherwise, since *T*_1_ ⊊ *S* and *S* is permanent, Theorem implies 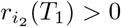 for some *i*_2_ ∈ *S*, and Proposition A.1 implies that there exists *T*_2_ ⊆ *S* such that 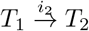 . If *T*_2_ = *S* we are done. Continuing inductively, at each step *n*, we obtain a path 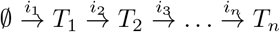 where *T*_1_, …, *T*_*n*_ are subsets of *S*. As the graph is acyclic, this induction stops for *n* less than or equal to the number of vertices in the community assembly graph restricted to *S*. As the system is permanent, Theorem 3 implies that *T*_*n*_ = *S*. Each edge of this path is a single-species transition between subcommunities of *S* in the graph produced by Step 3, so *S* is not removed in Step 4. If Step 5 prunes no edge between subcommunities of *S*, the path, and with it *S*, remains in the community assembly graph.

Now assume that all the vertices in the community assembly graph are permanent. We prove by induction on *k* = |*S*| that there is a directed path of length |*S*| from ∅ to *S*. If *k* = 1, then *S* = {*j*} for some *j*, and the path constructed above is 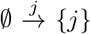. Now assume that the result holds for all communities with *k* ≥ 1 species satisfying the hypotheses, and let |*S*| = *k* + 1. Consider the last edge 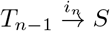 of the path constructed above. Since *T*_*n*−1_ ⊆ *S, i*_*n*_ ∈ *S, S* ⊆ *T*_*n*−1_ ∪ {*i*_*n*_} and *i*_*n*_ ∈*/ T*_*n*−1_, we have *S* = *T*_*n*−1_ ∪ {*i*_*n*_} and |*T*_*n*−1_| = *k. T*_*n*−1_ is permanent by assumption and the invasion graph restricted to *T*_*n*−1_ is acyclic. By the inductive hypothesis, there is a directed path of length *k* from ∅ to *T*_*n*−1_, and extending it by 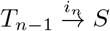 gives a directed path of length *k* + 1 = |*S*| from ∅ to *S*. □

### Proposition A.5.

*Let S be an end state of the community assembly graph corresponding to a permanent community. Then there exists m >* 0 *such that*

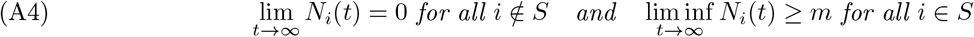

*whenever N*_*i*_(0) *>* 0 *for i* ∈ *S and N*_*i*_(0) *is sufficiently small for i* ∈*/ S. Moreover, if the invasion graph is acyclic and S*_1_, …, *S*_*n*_, *each corresponding to a permanent community, are the only communities of the invasion graph that cannot be invaded by any missing species, then for every initial condition with N*_*i*_(0) *>* 0 *for all i, (A4) holds with S* = *S*_*j*_ *for some* 1 ≤ *j* ≤ *n*.

*Proof*. The first assertion follows from the time average property of Lotka-Volterra systems [Hofbauer and Sigmund, 1998, Theorem 5.2.3] and Schreiber [2025, Proposition 4.6].

Now assume that the invasion graph is acyclic and *S*_1_, …, *S*_*n*_ are the only communities that cannot be invaded by any missing species. If some *S*_*j*_ = {1, …, *k*} for some *j*, then permanence implies that (A4) holds with *S* = *S*_*j*_ for all positive initial conditions. Otherwise, there is no equilibrium in (0, ∞)^*k*^, since {1, …, *k*} would then be a community that no missing species can invade. Let *E* = {*N* : Π_*i*_ *N*_*i*_ = 0} denote the extinction set. [Hofbauer and Sigmund, 1998, Theorem 5.2.1] implies *ω*(*N* (0)) ⊂ *E* for all *N* (0) ∈ (0, ∞)^*k*^. Since the invasion graph is acyclic, [Hofbauer and Schreiber, 2022, Lemma 4] implies that there are disjoint compact sets *K*_*j*_ ⊂ {*N* : *N*_*i*_ *>* 0 iff *i* ∈ *S*_*j*_} such that *ω*(*N* (0)) ⊂ *K*_1_ ∪ …∪ *K*_*n*_ for all *N* (0) ∈ (0, ∞)^*k*^. As *ω*(*N* (0)) is connected, *ω*(*N* (0)) ⊂ *K*_*j*_ for a single *j*. By compactness of *K*_1_, …, *K*_*n*_, there exists *m >* 0 such that (A4) holds with *S* = *S*_*j*_. □

## Numerical Simulations of Community Assembly

To simulate the assembly dynamics of the Lotka-Volterra model associated with a sequence of species introductions {*I*_1_, *I*_2_, …, *I*_*l*_} with 1 ≤ *I*_*j*_ ≤ *k*, we introduce each new species at a low density *ε >* 0, simulate the dynamics for a period of time *τ >* 0, remove any species whose density falls below *ε* at time *τ*, and then introduce the next species. Algorithm 3 describes this procedure in a formal way. Figures 1 and 7 provide examples of these simulations.

### Algorithm 3.

Assembly dynamics simulation for a Lotka-Volterra model

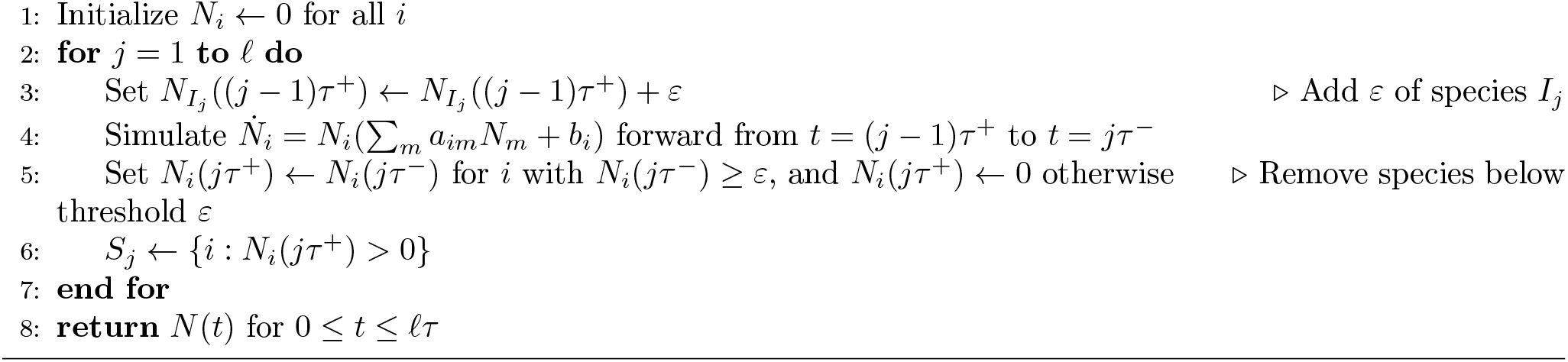

The following theorem shows that for *ε >* 0 sufficiently small and *τ >* 0 sufficiently large (depending on *ε*), the simulated assembly sequence corresponds to a valid path in the CAG. The theorem applies to models for which the CAG pipeline has been completed successfully, i.e., all vertices correspond to permanent communities and all directed edges have been verified to be dynamically realized.

### Theorem 4.

*Consider a Lotka-Volterra model for which all vertices of the community assembly graph correspond to permanent communities and all of its directed edges are dynamically realized. Let S*_0_ = ∅, *let* 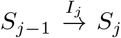,1 ≤ *j* ≤ *l, be the edges of a directed path in the community assembly graph, and let N* (*t*) *be the dynamics produced by Algorithm 3 for the introduction sequence* {*I*_1_, *I*_2_, …, *I*_*l*_}. *Then there exists ε*_0_ *>* 0 *and a function θ* : (0, *ε*) → (0, ∞) *such that such that, for all τ* ≥ *θ*(*ε*) *and* 1 ≤ *j* ≤ *l*, 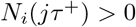 *if and only if i* ∈ *S*_*j*_.

*Proof*. For each vertex *S*≠ ∅ of the community assembly graph, let *A*(*S*) denote the global attractor of the Lotka-Volterra model restricted to *C*_+_(*S*) = {*N* : *N*_*i*_ *>* 0 iff *i* ∈ *S*}. *A*(*S*) is compact since *S* is permanent. Let *A*(∅) = {0}. Define N (*A*(*S*), *ε*) = {*N* : *N*_*j*_ = 0 for *j* ∈*/ S*, dist(*N, A*(*S*)) ≤ *ε*} to be the *ε* neighborhood of *A*(*S*). Let *ρ* be the minimum of *N*_*j*_ over *N* ∈ *A*(*S*), *j* ∈ *S*, and nonempty vertices *S* of the community assembly graph. *ρ >* 0 since each *A*(*S*) is a compact subset of *C*_+_(*S*) and there are finitely many vertices. For each directed edge 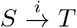, let 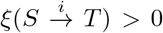 be given by the definition of dynamical realization (see Step 5 of the pipeline) with *η* = *ρ/*2. Set 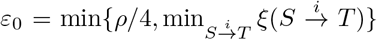. Fix *ε* ∈ (0, *ε* ). For each directed edge 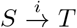 in the CAG, define 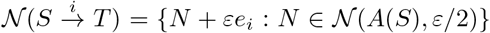, where *e*_*i*_ is the *i*-th unit vector. This set is compact and its points have resident densities at least *ρ* − *ε/*2 ≥ *η* and invader density 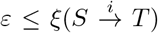, so dynamical realization implies that every solution starting in 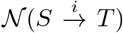 has its *ω*-limit set in *A*(*T* ). Since *r*_*j*_(*T* ) *<* 0 for all *j* ∈ (*S* ∪ {*i*}) \ *T*, Proposition A.5 implies that *A*(*T* ) is an attractor for the model restricted to {*N* : *N*_*j*_ = 0 for *j* ∈*/ S* ∪ {*i*}}. Hence, by compactness, there is a time 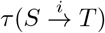 such that every solution starting in 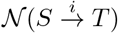 is within distance *ε/*2 of *A*(*T* ) for all 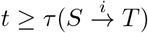. Set 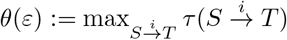.

Fix *τ* ≥ *θ*(*ε*). To complete the proof of the theorem, we show by induction that 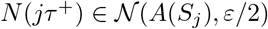 for 0 ≤ *j* ≤ *l*. For *j* = 0, *N* (0^+^) = 0. Now assume *N* ((*j* − 1)*τ* ^+^) ∈ *N*(*A*(*S*_*j*−1_), *ε/*2). Because *I*_*j*_ ∈*/ S*_*j*−1_, adding *ε* species *I*_*j*_ to this state, as Algorithm 3 does, leads to a point of 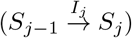, *N* (*jτ* ^−^) is therefore within *ε/*2 of *A*(*S*_*j*_). Species in *S*_*j*_ then have densities at least *ρ* − *ε/*2 *> ε* and all other species have densities at most *ε/*2 *< ε*. Hence, the removal step of Algorithm 3 sets the densities of species not in *S*_*j*_ to zero. As the densities of species not in *S*_*j*_ are zero in *A*(*S*_*j*_), this removal does not increase the distance to *A*(*S*_*j*_), and *N* (*jτ* ^+^) ∈ *N* (*A*(*S*_*j*_), *ε/*2). □

## Supplement B. Verifying permanence for cyclic invasion graphs

When the invasion graph restricted to a community *S* is cyclic, Theorem 3 does not establish permanence. In these cases, we use the criterion of Hofbauer [1981] (see also Hofbauer and Sigmund [1998, Theorem 13.6.1]).

### Theorem 5

(Hofbauer [1981]). *Assume the Lotka-Volterra model restricted to S is uniformly bounded. If there exist weights p*_*i*_ *>* 0, *i* ∈ *S, such that*

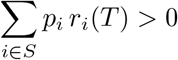

*for every proper subset T* ⊊ *S supporting an equilibrium x*^*^(*T* ), *then S is permanent*.

Following Jansen [1987], the existence of such weights can be decided by linear programming: maximize *ε* subject to ∑_*i*∈*S*_ *p*_*i*_*r*_*i*_(*T* ) ≥ *ε* for all such *T, p*_*i*_ ≥ 0 for all *i* ∈ *S*, ∑_*i*∈*S*_ *p*_*i*_ = 1, and *ε* ≤ 1. Permanence is established if the optimal *ε* is positive. The criterion is sufficient but not necessary (see Discussion).

## Supplement C. Parameterizations for the Empirical Examples

### The New Zealand Grassland. Dormann [2008, equation (1)] used a Lotka-Volterra parameterization of the form

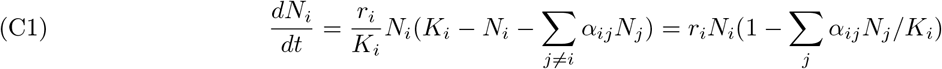

Where

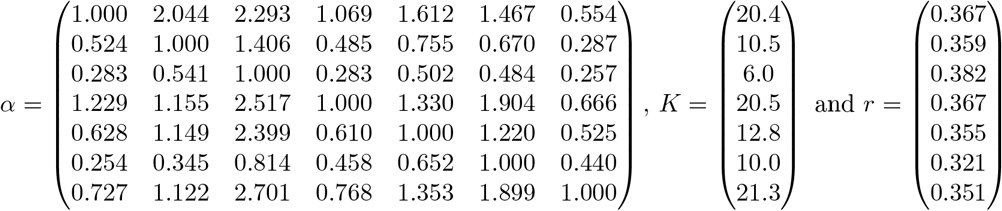

Defining *b*_*i*_ = *r*_*i*_ and *a*_*ij*_ = −*r*_*i*_*α*_*ij*_*/K*_*i*_ gives our formulation of the Lotka-Volterra model

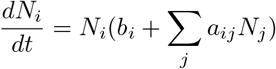

Where

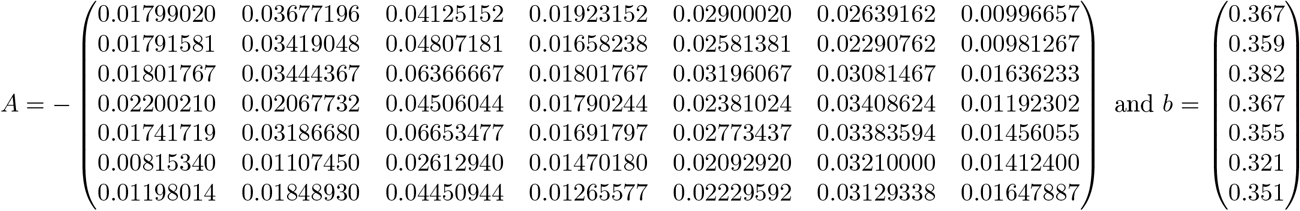

### The European Pasture

Geijzendorffer et al. [2011, equation (5)] parameterized a Lotka-Volterra model of the form

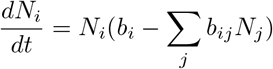

which corresponds to our formulations of the Lotka-Volterra model with *a*_*ij*_ = −*b*_*ij*_. The parameters for the spring are given by

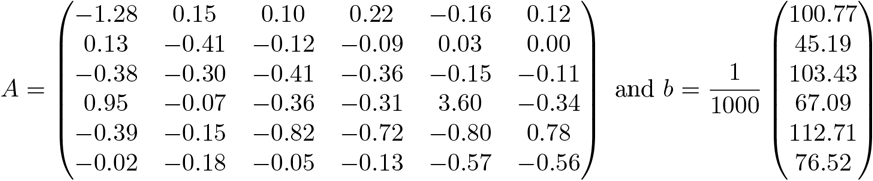

## Supplement d. example of indirect blow-up

Here, we provide an empirical Lotka-Volterra model for which all species pairs satisfy the *B*-matrix condition, but the three species community does not (Figure D0). The interaction and intrinsic growth matrices come from [Spaak et al., 2021] based on data by Caputa [1948]:

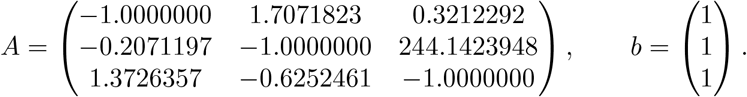

**Figure D0.**
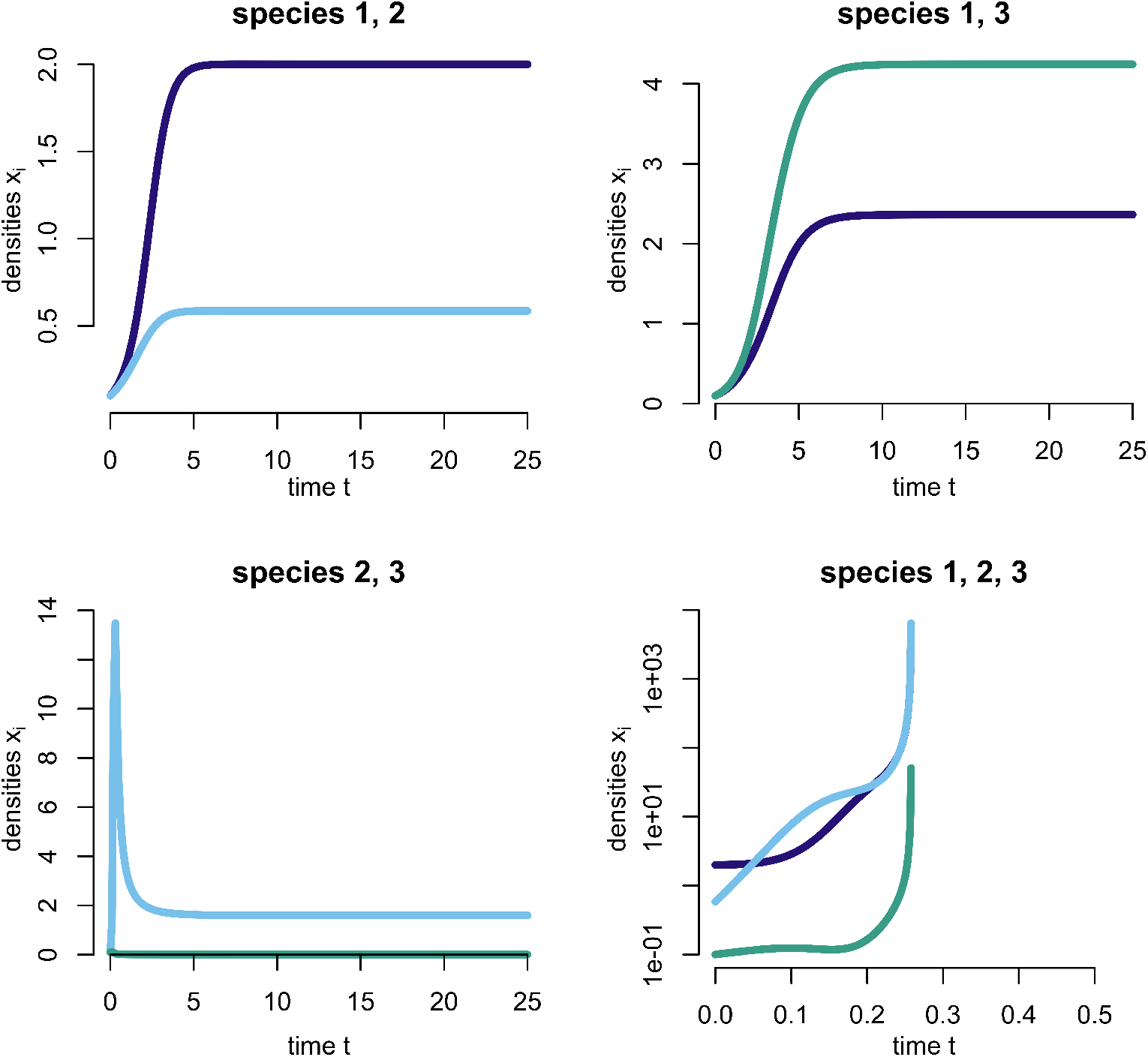
Boundedness can fail without warning from subcommunities. All three pairs (top-left, top-right, bottom-left) approach bounded states, whereas the full triad (bottom-right, log scale) diverges.

